# Lower Brain Glucose Metabolism in Normal Ageing is Predominantly Frontal and Temporal: A Systematic Review and Pooled Effect Size and Activation Likelihood Estimates Meta-Analyses

**DOI:** 10.1101/2022.08.08.503243

**Authors:** H.A Deery, R Di Paolo, C. Moran, G.F. Egan, S.D. Jamadar

## Abstract

This review provides a qualitative and quantitative analysis of cerebral glucose in ageing. We undertook a systematic review of the literature followed by pooled effect size and Activation Likelihood Estimates (ALE) meta-analyses. Studies were retrieved from PubMed following the PRISMA guidelines. After reviewing 653 records, 22 studies with 24 samples (n = 993 participants) were included in the pooled effect size analyses. Eight studies with 11 samples (n = 713 participants) were included in the ALE analyses. Pooled effect sizes showed significantly lower cerebral metabolic rates of glucose for older versus younger adults for the whole brain, as well as for the frontal, temporal, parietal and occipital lobes. Among the sub-cortical structures, the caudate showed a lower metabolic rate among older adults. In sub-group analyses controlling for changes in brain volume or partial volume effects, the lower glucose metabolism among older adults in the frontal lobe remained significant, whereas confidence intervals crossed zero for the other lobes and structures. The ALE identified nine clusters of lower glucose metabolism among older adults, ranging from 200mm^3^ to 2,640mm^3^. The two largest clusters were in the left and right inferior frontal and superior temporal gyri and the insula. Clusters were also found in the inferior temporal junction, the anterior cingulate and caudate. Taken together, the results of the meta-analyses are consistent with research showing less efficient glucose metabolism in the ageing brain. The findings are discussed in the context of theories of cognitive ageing and are compared to those found in neurodegenerative disease.

## 1. Introduction

### 1.1 The brain requires a highly responsive and dynamic supply of glucose

Although the brain cannot store glucose, it relies on glucose as its primary energy source. The adult brain accounts for approximately 2% of total body weight but requires around 20% of total glucose supply (Kety, 1957; Sokoloff, 1960). The basal metabolic demand of the brain is estimated to be 30% of whole-brain glucose use, providing the fuel required for functions such as resting state connections and neuronal action potentials (Hyder, 2013; Tomasi et al., 2013). Seventy percent of brain glucose use is for spontaneous functional processes, such as signal processing and cortical computation, including the production of action potentials, postsynaptic potentials, and the maintenance of ion gradients (Mergenthaler et al., 2013; Tomasi et al., 2013).

There are complex and dynamic changes in energy consumption of brain cells from moment to moment, as the brain performs its vast array of functions (Yellen, 2018). Brain glucose metabolism is tightly coupled to cerebral blood flow, to the degree that local blood flow is highest in regions with the highest glucose metabolism (Clarke & Sokoloff, 1999). Functional brain activity is accompanied by an increase in cerebral blood flow and in glucose uptake to meet the acute energy demands of neurons (Diaz-Garcia & Yellen, 2019). The dynamic demands for neural glucose require a highly responsive system that is controlled by both feedback and feedforward processes, and that produces changes in flow through specific biochemical pathways (Yellen, 2018). To support this highly responsive system, an intricate interplay exists between the brain, endocrine system, central and peripheral energy supply, and energy utilisation (see Cunnane et al, 2020; Camondola & Mattson, 2017 for reviews of these mechanisms). The level of cerebral glucose metabolism is considered a reliable measure of neuronal activity (Mergenthaler et al., 2013). Here, we present a systematic review, effect size meta-analysis and coordinate-based meta-analysis to examine the convergence of evidence for cerebral metabolic glucose differences across the adult lifespan.

### 1.2 Assessing cerebral glucose metabolism changes in neurodegeneration and ageing

Developed over 40 years ago, F-18 fluorodeoxyglucose (^18^F-FDG or FDG) positron emission tomography (PET) is the most commonly used neuroimaging technique for directly evaluating brain metabolism *in vivo* (Huang et al., 1980). FDG-PET has been widely used to study alterations in cerebral glucose in ageing and disease states (see Mielke et al.,1988; Stranahan & Mattson, 2012). FDG is metabolically trapped in neurons or glial cells after being metabolised into glucose-6-phosphate. The local concentrations of the tracer radioactivity can be measured using FDG-PET. Quantitative measures of FDG-PET require arterial blood sampling at multiple time points. A mathematical model that describes the kinetics of FDG transport is used to determine the transport rate constants of FDG and to convert the radioactivity measurements to metabolic rates (Huang et al. 1980). For a review of tracer kinetic modelling for PET, see Carson (2003).

Quantitative measures of FDG uptake have shown that brain glucose metabolism changes across the lifespan, and that there are archetypical regional patterns of hypometabolism, particularly in neurodegenerative diseases (see Chen & Zhong, 2013; Butterfield & Halliwell 2019; Cunnane et al., 2020). In mild cognitive impairment and Alzheimer’s disease, regions of the default mode network and the medial temporal lobe are most affected early in the disease process. Meta-analyses comparing Alzheimer’s patients to controls show that show the greatest reduction in glucose metabolism include the precuneus and temporal, supramarginal, cingulate, fusiform, angular, inferior parietal and middle frontal gyri (He et al., 2015; also see Mosconi, 2013, for a review). They also include the left precentral and parahippocampal gyri and right superior frontal gyrus and thalamus (He et al., 2015).

### 1.3 Cerebral glucose metabolism in “normal” ageing

#### 1.3.1 Macro-Scale Changes

The results of FDG-PET studies on CMR_GLC_ in ageing have often been reported at the whole-brain or lobular level, and have been inconsistent. Some early studies showed no significant difference in CMR_GLC_ among older adults compared to younger adults (Duara er al., 1984; De Leon et al., 1984); whereas others have reported a significant decline, particularly in the frontal region (Alavi, 1989; Chawluk et al., 1987; Ivançević et al., 2000; Kuhl et al., 1982; Pardo et al. 2007; Petit-Taboué et al., 1998).

#### 1.3.2 The Need for Assessment of the Locations of Metabolic Changes in Ageing

An improved understanding of the location of cerebral glucose metabolism changes in ageing is an important empirical and clinical matter, as better characterisation of the timeframe and profile of normal ageing and neurodegeneration may help to prevent or slow cognitive decline and disease trajectories. Although it is informative to assess cerebral glucose metabolism in ageing at a macro-scale, voxel-based image analysis can improve accuracy by providing a more precise assessment of the locations of metabolic changes with age. Voxel-based quantitative analysis methods also provide statistical mapping of whole brain that can detect areas missed in analysis at the lobular level or in region of interest (ROI) analysis limited to specific brain areas.

#### 1.3.3 Concurrent Age-Related Brain Changes Impacting Brain Metabolism

The variability in findings of age differences in cerebral glucose metabolism may reflect concurrent age-related changes that occur in the brain, that can affect absolute CMR_GLC_ and have been accounted for in a limited number of studies. For example, the brain loses volume and cortical thickness with age and displays an expansion of the ventricular system, resulting in less metabolically active tissue (Liu et al., 2017). Grey matter volume undergoes a five-fold growth in the first decade of life (Courchesne et al. 2000), reaches its maximum in early adulthood and then shows a relatively linear reduction, with declines doubling between 30-40 and 70-80 years of age (Sigurdsson et al, 2012; Battaglini et al. 2018; Toepper, 2017; Zanto & Gazzley, 2019). Brain atrophy is also accompanied by increased cerebrospinal fluid spaces (Tanna et al., 1991). It is now accepted that age-related brain atrophy reduces CMR_GLC_ calculated on a voxel-wise basis, due to reduced metabolically active tissue and an increased fraction of non-metabolising cerebrospinal fluid (Tanna et al., 1991).

Multi-modal imaging has been adopted to allow for the adjustment of volume and cortical thickness losses with age when studying brain glucose metabolism. In early studies, PET images were co-registered to CT scans (de Leon et al.,1987) and ventricle size was measured and used to adjust for brain volume loss in ageing (Kushner et al., 1987). More recently, PET images have been co-registered to each participant’s MR image of grey matter with non-brain tissue and cerebrospinal fluid removed from the images during processing (Nugent et al, 2014a, b).

The *partial volume effect* (PVE) is a phenomenon that reduces the quantitative accuracy of PET images. In a PET image, the intensity of a particular volume reflects the tracer concentration not only of the tissue within that volume but also in the surrounding area (Yang et al., 2017). This means that the measured tracer activity concentrations are not accurate as the signals spill over into surrounding volumes. The partial volume effect becomes problematic when the dimensions of a region of interest are less than two to three times the full width at half maximum spatial resolution of the PET scanner (Hoffman et al., 1979). The partial volume effect is also particularly marked when cortical atrophy is present, such as in the ageing brain (Meltzer et al., 1999).

PET scanners used in the neural metabolic ageing literature in 1980s to the early 2000s typically had spatial resolution (full width at half maximum) of 6-17mm (e.g., de Leon et al., 1983), whereas recent scanners provide resolution in the 2-4mm range (e.g., Nugent, 2014a, b). These spatial resolution values mean that partial volume effects are likely present in brain regions smaller than approximately 18-50mm in neural metabolic ageing studies conducted prior to early 2000s, and in regions smaller than 6-12mm in more recent studies. Researchers have often parcellated the brain into 40 or more regions of interest, creating volumes at the sizes or smaller where partial volume effects are likely impacting the results (see Table 1 and 2 for parcellation approaches and region of interests used in the literature).

**Table 1.**
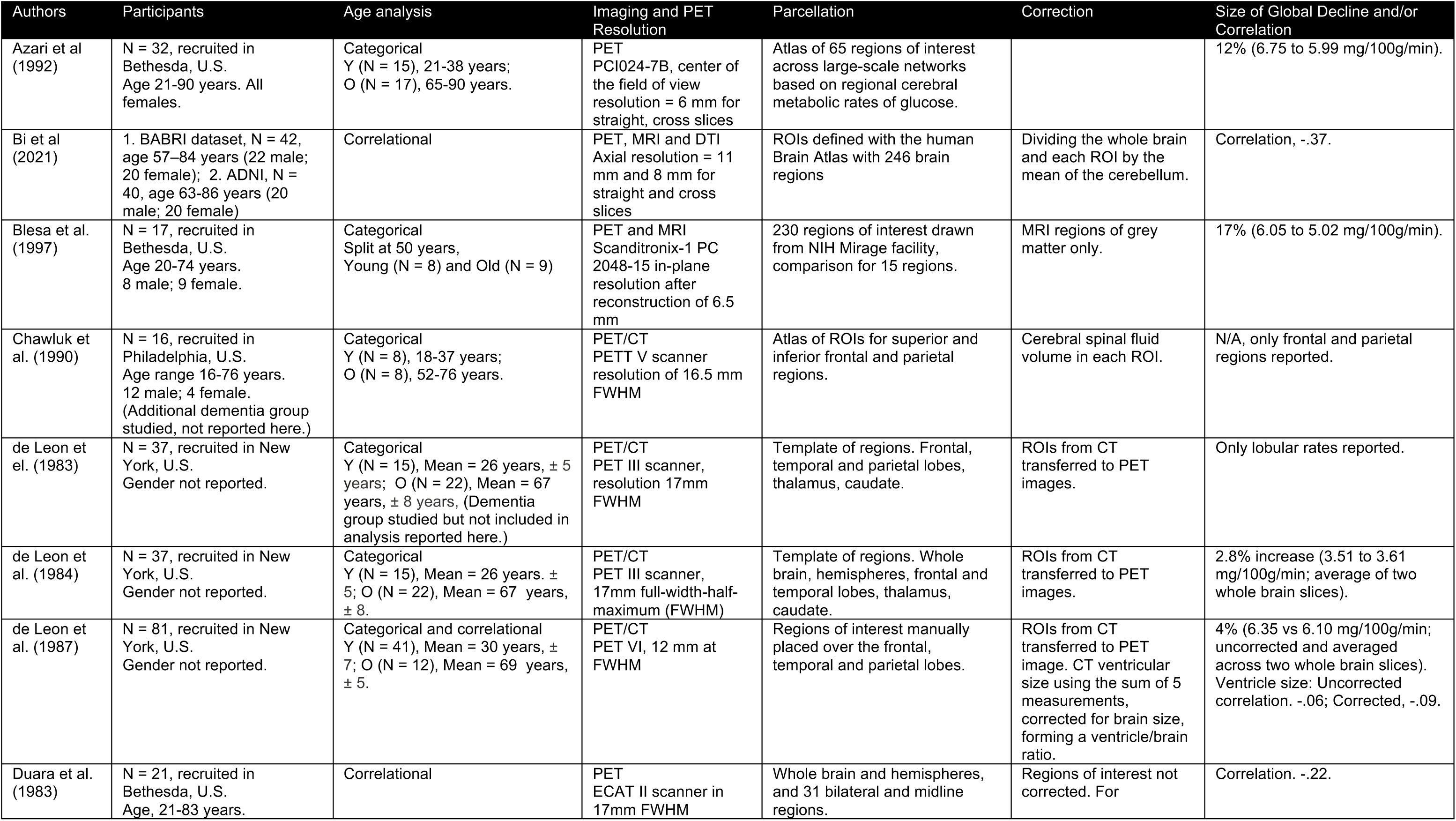
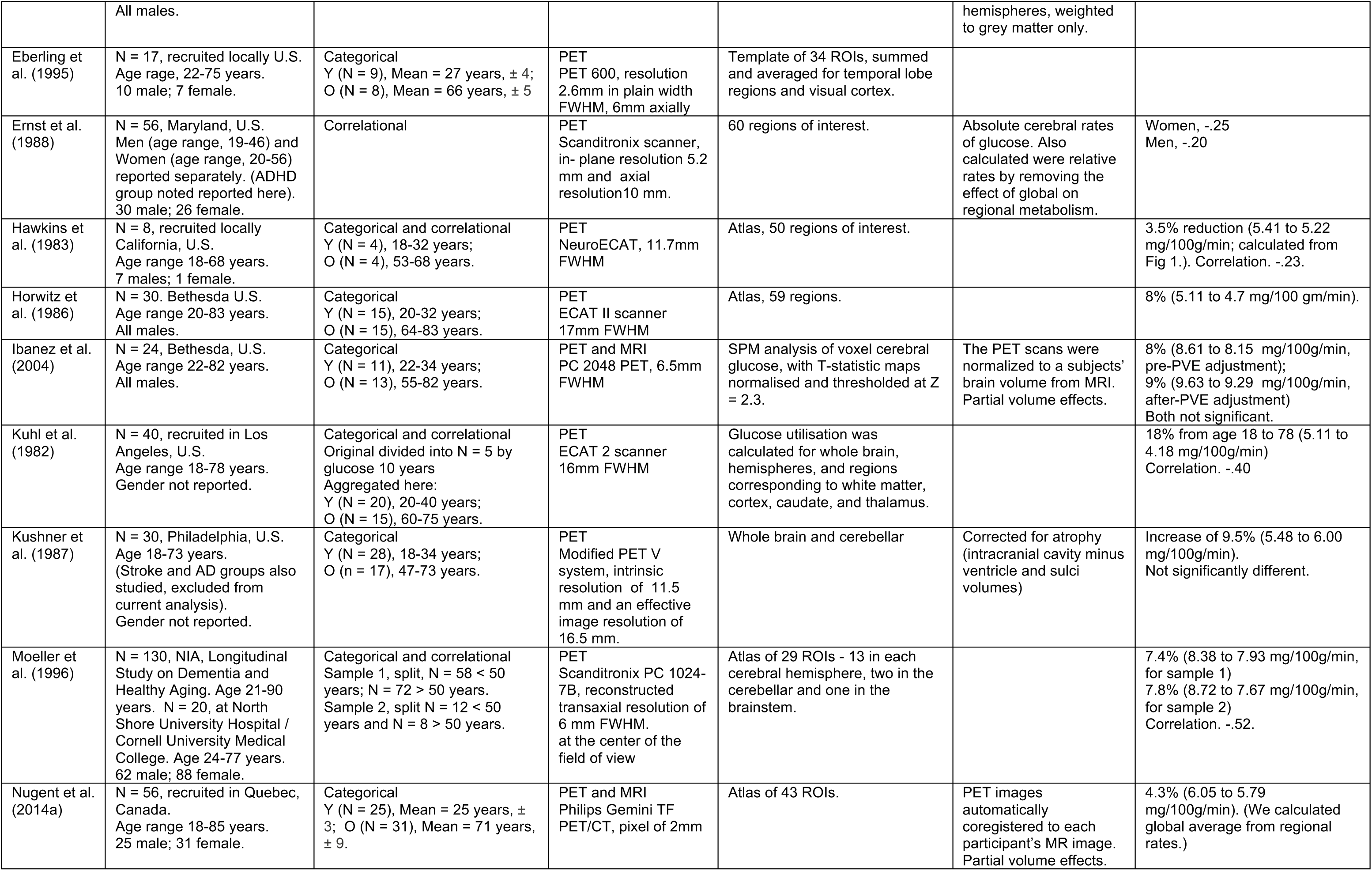
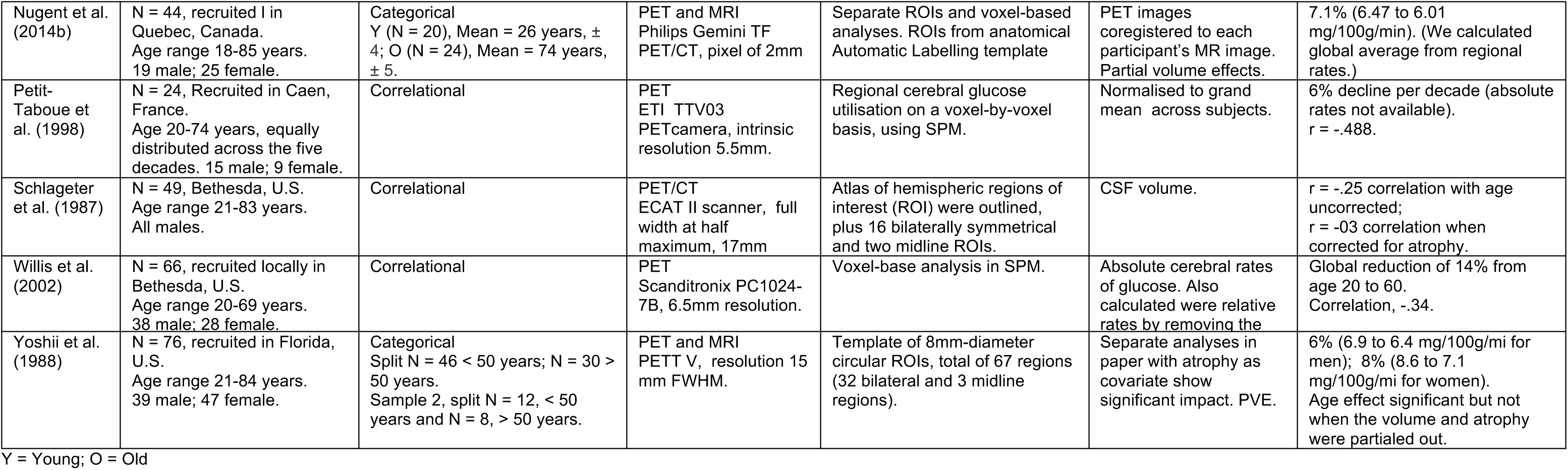
Studies included in the effect size meta-analyses of the cerebral rate of glucose across the adult lifespan.

**Table 2.** Studies included in ALE meta-analysis

| Authors | Sample # | Participants | Reported Coordinates | Imaging | Analysis | Correction |
| --- | --- | --- | --- | --- | --- | --- |
| 1. Ibanez et al. (2004) | 1. All | N = 24, recruited in Bethesda, U.S.<br>Young, 22-34; Old, 55-82 years. All male. | Local maxima of brain regions in which mean rCMR<sub>glc</sub> was significantly less in older than in young subjects. | PET and MRI | SPM analysis of voxel cerebral glucose, with T-statistic maps normalised and thresholded at Z = 2.3. | Partial volume effects. |
| 2. Kalpouzos et al. (2009) | 2. All | N = 45, recruited in Caen, France.<br>Y = young 20-38; Middle-aged, 40-58; Older, 60-83 years.<br>21 male; 24 female. | Regions of glucose metabolism decreasing with age, local maximum activity of clusters. | PET and MRI | SPM voxel-level analysis of PET FDG uptake and MRI. | Partial volume effects and coregistration to MR data. |
| 3. Kim et al. (2009) | 3. Males<br>4. Females | N = 78, recruited locally, Busan, South Korea.<br>Age continuous, 15-81 years.<br>32 male; 46 female. | Age-associated decreased uptake of FDG in grey matter for males and females separately. | PET | Voxel-based analysis. |  |
| 4. Pardo et al. (2007) | 5. All | N = 46, recruited locally, University of Minnesota, U.S.<br>Age continuous, 18-90 years.<br>22 male; 24 female. | Local maxima of clusters with negative correlations of glucose with age. | PET | Voxel-based analysis and correlation between glucose uptake and age. |  |
| 5. Petit-Taboue et al. (1998) | 6. All | N = 24, Recruited in Caen, France.<br>Age range, 20-74 years.<br>15 male; 9 female. | Peaks of most significant declines in CMR<sub>glc</sub> with age (P < 0.001). CRM<sub>GLC</sub> normalised to grand mean across subjects. | PET | Cerebral glucose utilisation on a voxel-by-voxel basis, using SPM. |  |
| 6. Shen et al. (2012) | 7. Males<br>8. Females | N = 234, recruited in Hangzhou, China<br>Age continuous, 26 - 77 years<br>126 male; 108 female. | Regions of glucose metabolism decreases with aging in the male and female participants separately. | PET | Voxel-based analysis of cerebral rate of glucose using a general liner model (GLM). |  |
| 7. Yanase et al. (2005) | 9. Males<br>10. Females | N = 139, recruited locally in Kanazawa, Japan<br>Age continuous, 24-81 years<br>71 male; 68 female. | Peaks of significant decreases in relative FDG activity with age in men and women separately before and after PVE correction (PVE corrected data used here). | PET and MRI | SPM, voxel-based analysis. | Partial volume effects and coregistration to MR data. |
| 8. Yoshizawa et al. (2014) | 11. All | N = 123, recruited locally in Tokyo, Japan.<br>Age continuous, 33-78 years.<br>91 male; 73 female. | Clusters of negative correlation between age and FDG uptake. | PET | Statistical parametric mapping of FDG signal. |  |

Several methods are available for the correction of partial volume effects in PET imaging (for a review, see Frouin et al. 2002; Meechai et al., 2014). Some PVE correction techniques use MR images to define the boundaries of the region of interest based on anatomical *a priori* information. Several algorithms have also been developed to correct for partial volume effects (Yang et al., 2017) and have been applied to PET studies of metabolic brain changes in ageing (e.g., Ibáñez, 2004). These algorithms typically use a pre-calculated correction factor, based on the relationship between the volume diameter and the ratio of adjusted activity and the true activity (Meehcai et al., 2014).

The literature on whether or not brain volume changes and PVE can account for brain glucose metabolism reductions in ageing is inconclusive. There is evidence that age-related whole brain, frontal, parietal, and temporal reductions in the absolute CRM_GLC_ are not significant after controlling for volume and brain atrophy (Schlageter et al., 1987; Yoshii et al., 1988). Some authors (e.g., Curiati et al., 2011; Ibáñez et al., 2004; Yanase et al. 2005) have reported reductions in CMR_GLC_ among older adults in several cortical regions and subcortical structures. However, CMR_GLC_ reductions were not significant after adjusting for partial volume effects. Others have found regional differences in age-related CMR_GLC_ decline, even after controlling for partial volume effects (e.g., Kalpouzos et al., 2009; Nugent et al., 2014a, b). For example, Nugent et al. (2014) found lower glucose metabolism among older adults in the frontal lobe, temporal cortex, anterior cingulate, insula, putamen and thalamus after removing non-brain tissue and cerebral spinal fluid and controlling for partial volume effects.

In reviewing the literature on age-related changes in CMR_GLC,_ it is apparent that the sample sizes and associated effect sizes vary across studies. Some studies are limited to sample sizes less than 10 and report whole-brain CMR_GLC_ reductions as little as 3.5% (e.g., Hawkins, 1983), whereas others have included samples larger than 40 (Yoshi et al., 1988) or even 100 (Moeller et al., 1996) and reported CMR_GLC_ reductions as high as 18% (Kuhl et al., 1992). The differences in study sample and effect sizes mean that the relative contribution of these studies to the body of evidence on age-related differences varies and needs to be evaluated. However, to our knowledge, the literature has not been systematically reviewed and meta-analysed. Meta-analysis offers the benefit of pooling the effects across studies, giving effect sizes of studies with a higher precision (i.e., a smaller standard error), a greater weight in the analysis.

In summary, the literature reviewed above indicates that brain glucose metabolism changes across the adult lifespan and that disruptions to normal glucose homeostasis impact brain function and cognition. However, an understanding of the effect size of age-related changes in “normal” ageing at a whole-brain and regional level is currently limited. Moreover, there is a need to understand whether age differences in CMR_GLC_ persist after brain volume changes and PVE are taken into account. An improved understanding of the location of cerebral glucose metabolism changes in ageing is also an important empirical and clinical matter. Such an understanding is important as evidence suggests that the location and time course of alterations to brain glucose metabolism is different in normal ageing and dementia. An improved understanding of these issues is especially important in ageing and neurodegeneration, as changes can begin to occur several decades before the onset of disease or symptoms (Jack et al., 2010). Better characterisation of the timeframe and profile of normal ageing, cognitive decline and neurodegeneration may represent an opportunity to delay, slow or even reverse or prevent neurodegenerative processes and improve the quality of life of older adults.

### 1.4 Scope of the Current Review

We combine a systematic review of the literature and meta-analyses to provide a qualitative and quantitative account of cerebral metabolic glucose differences across the adult lifespan. We adopt the PRISMA method to systematically review the literature on studies of metabolism in ageing and use two complementary meta-analyses techniques to combine the results from across the studies based on the nature of the data reported. First, pooled effect size meta-analyses are used to assess the global and regional CMR_GLC_ differences in ageing. We assess and quantify age differences for younger and older adults at the whole brain and lobular level, as well as for white matter and sub-cortical structures. Second, we use Activation Likelihood Estimation Meta-analysis (ALE) to examine the regional convergence of age-related differences in CMR_GLC_. ALE is an objective, quantitative technique for coordinate-based meta-analysis of neuroimaging results that has been validated for a variety of uses, including to compare brain activity across populations, such as older and younger adults (Turkeltaub et al., 2012).

## 2 Method

### 2.1 Type of Studies and Participants

Studies of CMR_GLC_ across the adult lifespan were retrieved from PubMed in June 2021, following the PRISMA 2020 statement (Page et al., 2021). The identification, screening and selection process is summarised in Figure 1. We included studies in which participants were adults aged between 18 and 90 years of age and in which the mean and standard deviation of CMR_GLC_ were reported for older and younger adults, or the correlation with age and CMR_GLC_ across the adult lifespan was reported. Studies that reported age differences in the CMR_GLC_ for the whole brain, lobes and/or sub-cortical structures were included. We also included studies in which whole-brain analyses were undertaken of CRM_GLC_ and the significant age-group activation differences (either higher or lower in older versus young adults) were reported for use in the ALE meta-analyses.

**Figure 1.**
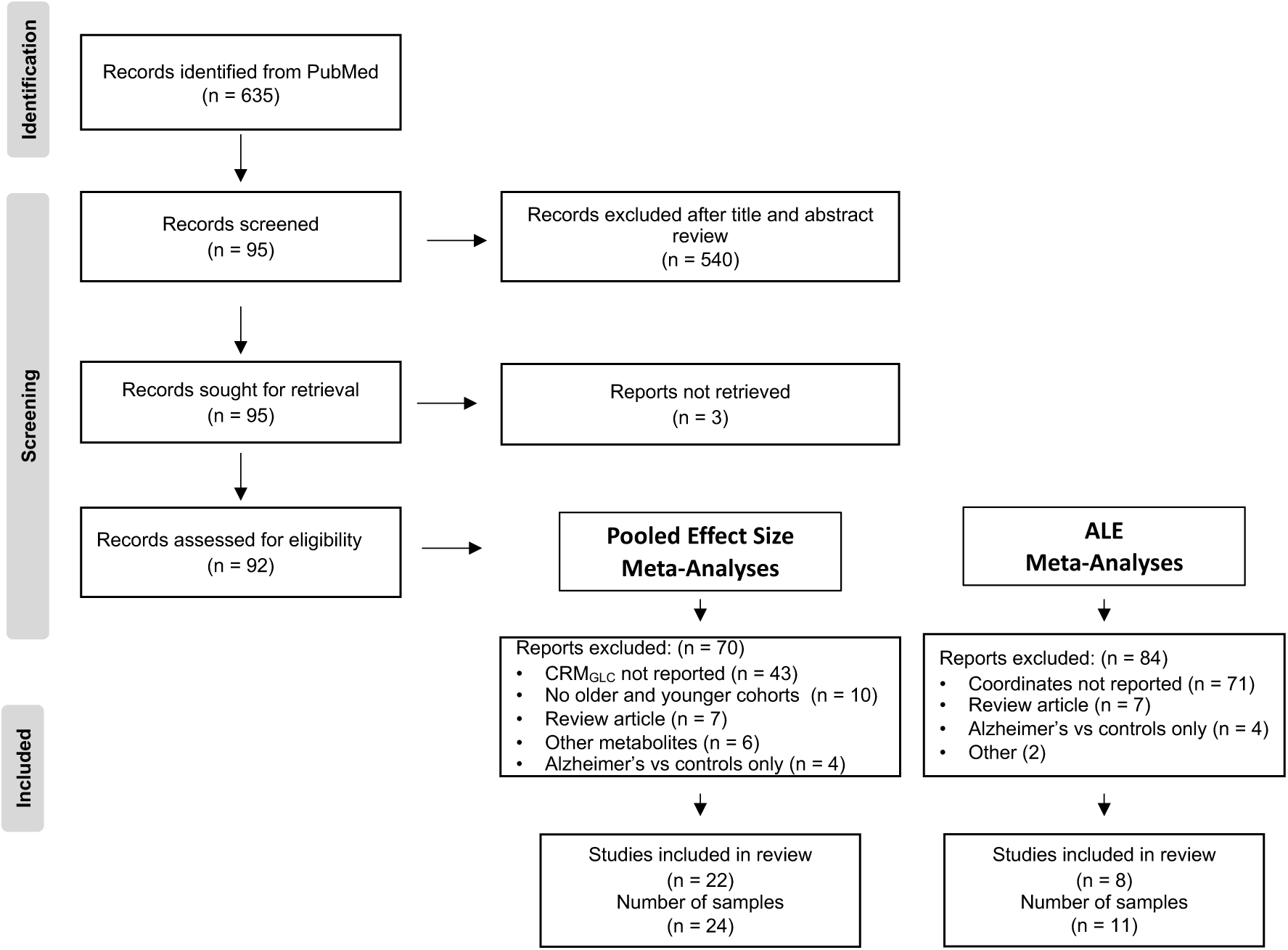
The identification, screening and selection process of studies included in the systematic review and meta-analyses.

**Figure 2.**
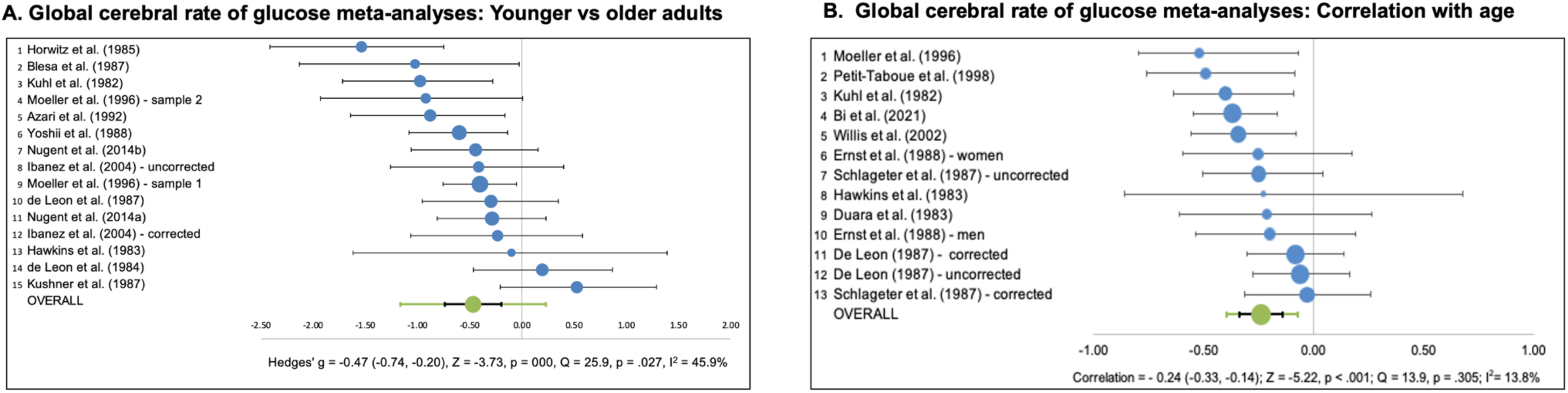
Global cerebral rate of glucose meta-analyses. A. Younger vs older adults; B. Correlation with age. For each study, the blue dot show Hedges’ g effect size on the X-axis and the horizontal line illustrates the high and low limits of the 95% confidence interval. The size of the blue dot reflects the weight of the study in the overall effect size in the bottom row. For the overall effect, Hedges’ g effect size (green dot), its confidence interval (black) and its prediction interval (green) are shown. The prediction interval gives the 95% confidence range in which the outcome of a future study will fall, assuming that the effect sizes are normally distributed.

Study participants were classified as “normal” or “healthy”. The study authors reported screening participants for medical history and undertook physical and neurologic examinations. Exclusion criteria included brain diseases, hypertension, cardiovascular or cerebrovascular disease, diabetes, substance abuse or addiction, head trauma, psychiatric illness, or systemic illnesses known to influence cognitive function.

### 2.2 Search Strategy

Search criteria were based on combinations of the following keywords and terms: brain glucose, brain glucose metabolism, cerebral rate of glucose, metabolic rate of glucose, cerebral glucose utilisation, regional cerebral metabolic rate of glucose, brain fuel, brain energy, FDG-PET and 18F-FDG. Additional papers were identified by manually searching the citations from the retrieved references. Studies were included if they were published in a peer-reviewed journal in English, and used human subjects.

### 2.3 Exclusion Criteria

For the pooled effect size meta-analyses, papers were excluded if they did not report the mean and standard deviation of the CMR_GLC_ for older and younger adults, nor a correlation with age. Papers that reported standardised uptake values (SUV) only were also excluded. Standard uptake values are calculated as the ratio of tissue activity concentration and administration dose, divided by body weight and are often normalised to the whole brain or a region (Nugent et al., 2020). Standard uptake values have the advantage of not requiring blood sampling or dynamic imaging. However, standard uptake values are semi-quantitative and do not allow for purely quantitative imaging of CMR_GLC_. Papers were also excluded if they did not report the use of a kinetic model to quantify CMR_GLC_. Disease states (e.g., metabolic dysfunction and mental disorders including developmental disorders, dementias, epilepsy) were excluded in the search criteria.

For the ALE meta-analyses, papers were excluded if they did not report significant peak age-group activation differences from whole-brain analysis; Studies were excluded if they reported region of interest analysis only (e.g., Curiati et al., 2011) in line with recommendations for ALE meta-analyses (Müller et al., 2018).

### 2.4 Selection Process and Included Studies

The literature search retrieved 635 unique records. Titles and abstracts were independently screened and reviewed by two raters (HD & RDP) for inclusion, which reduced the articles for retrieval and review against the inclusion criteria to 92. Where differences were found between the raters, the articles for inclusion were discussed and reconciled.

### 2.5 Meta-Analyses

#### 2.5.1 Pooled Effect Size Meta-Analyses

Data was entered into Meta-Essentials tool (https://www.erim.eur.nl/research-support/meta-essentials/) and analysed using a Random Effects Model for pooled effect sizes. Analyses were undertaken separately for studies reporting the global mean CMR_GLC_ for young versus older adults, as well as the correlation of age and CMR_GLC_. Analyses were also undertaken for the frontal, temporal, parietal and occipital lobes, for white matter and for available subcortical structures: caudate, cerebellum and thalamus.

Five studies reported CRM_GLC_ for regions within some or all lobes but did not provide overall lobe CRM_GLC_ values. We averaged the regions within the lobes to arrive at lobular CRM_GLC_ values. Where studies reported the CMR_GLC_ for the contralateral right and left regions separately (e.g., Azari et al., 1992; de Leon, 1983; Moeller, 1996), we also averaged those regions. For those studies reporting values in *μ*.mol per 100g^-1·min-1^, we converted the values to mg/100ml/min for use in the meta-analyses using the molar mass of FDG (multiplying by 181.1 g/mol and dividing by 1,000).

For the pooled effect size measure, *Hedge’s g* is reported. Hedge’s g is based on the difference of the means in units of the pooled standard deviation. *Cochran’s Q* was used as the test for heterogeneity of the effect size of the meta-analyses. Based on a chi-square distribution, it generates a probability that, when large, reflects substantial variation across studies rather than within subjects in a study. A statistically significant result was considered to reflect a need to investigate the underlying heterogeneity and that meta-analytical *sub-group analyses* may help identify the mediators of sub-group differences.

Where sufficient samples were available, meta-analytic sub-group analyses were undertaken to compare the effect sizes from studies that did and did not correct for brain volume, partial volume effects, or both. ANOVA was used to assess differences in sub-group effects relative to the precision of the difference. A p-value of less than 0.1 indicates a statistically significant sub-group effect (Richardson et el., 2019).

*I^2^* was used to quantify heterogeneity and calculates the proportion of variation due to heterogeneity rather than chance. Value ranges from 0% to 100%, with higher values indicating greater heterogeneity. Together with the p-value for the chi-square test: 0% to 40% was considered to be unimportant; 40% to 60% as representing moderate heterogeneity; and above 60% representing substantial heterogeneity (Higgins & Green, 2011).

As a large number of studies were conducted pre-2000’s on models of PET equipment with poor spatial resolution, we also conducted sub-group analyses for scanner resolution but found no significant effect (all p>.1). We also divided the studies pre-and post-2000 and by decade of publication and found no impact (all p>.1).

#### 2.5.2 ALE Meta-Analysis

Local foci coordinates were entered into GingerALE 3.0.2 (https://www.brainmap.org/ale/.) GingerALE computes the *activation likelihood estimation* value for each voxel in the brain, which is an estimation of the likelihood that at least one of the foci in a dataset is truly located at that voxel. GingerALE determines the null distribution of the ALE statistic at each voxel using Gaussian probability density distributions and a random effects model that tests for agreement across experimental groups and limits the effect of a single study. A conservative threshold of p < 0.001 was chosen for thresholding the ALE statistic maps to minimise Type I error.

Cluster analysis was performed to find the neighbouring volumes above the cluster threshold. We adhered to the recommendations of Eickhoff et al. (2012) by using a cluster-level family-wise error of p < 0.05 to correct for multiple comparisons, following an initial cluster forming threshold of uncorrected p < 0.001. Statistics for the clusters above the threshold are reported, including volume and the locations and values of the peaks within the cluster. Anatomical labels were included based on Talairach Daemon (talairach.org) for the peaks and volumetric label data for the clusters. Where a cluster crossed lobes or sulcus, to aid interpretation we investigated the composition of the clusters further by plotting the location of the activation from each contributing study (see Figures 3 and 4).

**Figure 3.**
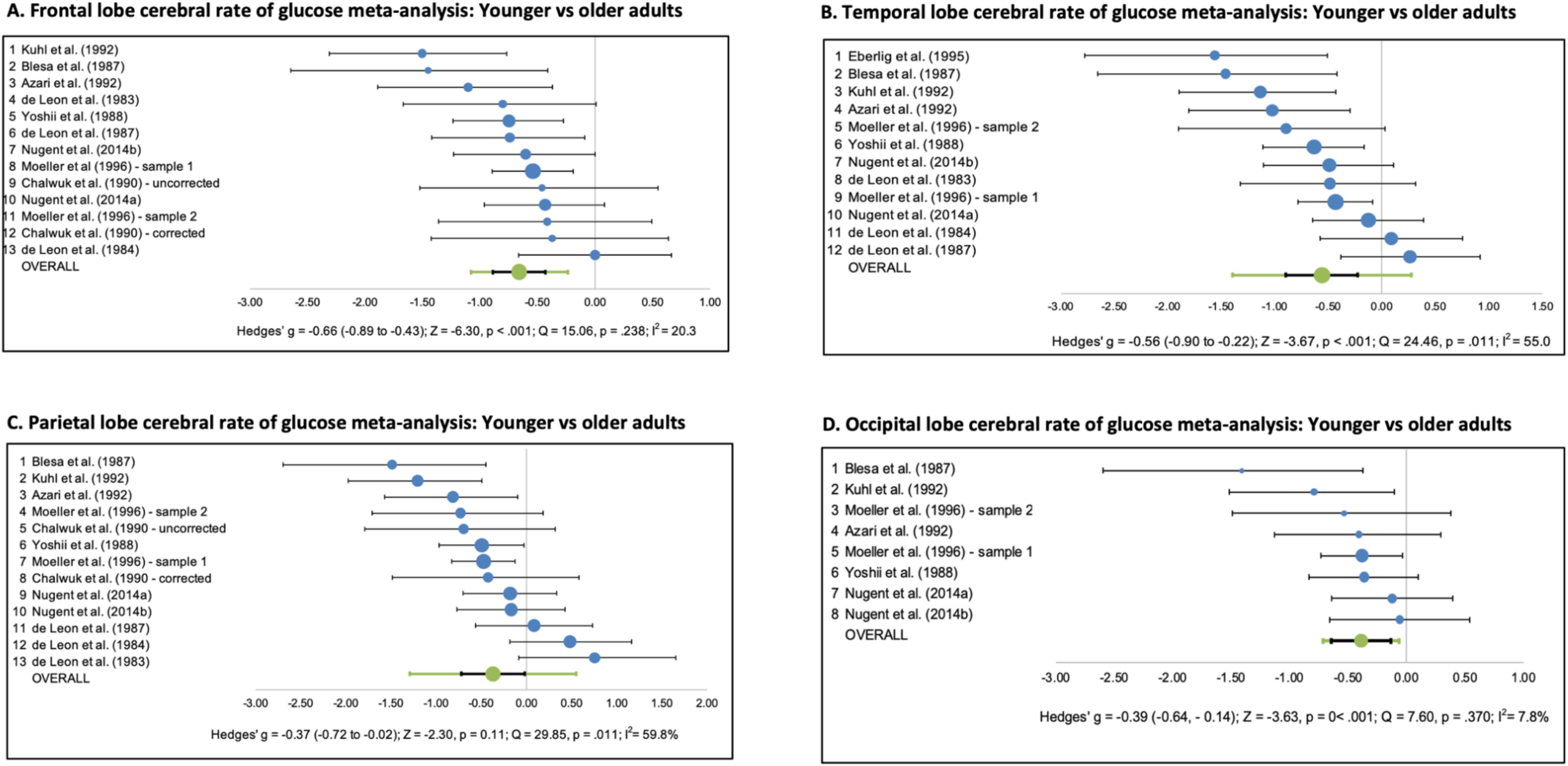
Lobular cerebral rate of glucose meta-analyses for younger vs older adults; A. Frontal lobe; B. Temporal lobe; C. Parietal lobe; D. Occipital lobe. For each study, the blue dot show Hedges’ g effect size on the X-axis and the size of the dot the weight of the study in the overall effect size. The horizontal line illustrates the high and low limits of the 95% confidence interval. The overall effect size is shown in the bottom row, with Hedges’ g effect size (green dot) and its confidence interval (black) and its prediction interval (green). The prediction interval gives the 95% confidence range in which the outcome of a future study will fall, assuming that the effect sizes are normally distributed.

**Figure 4.**
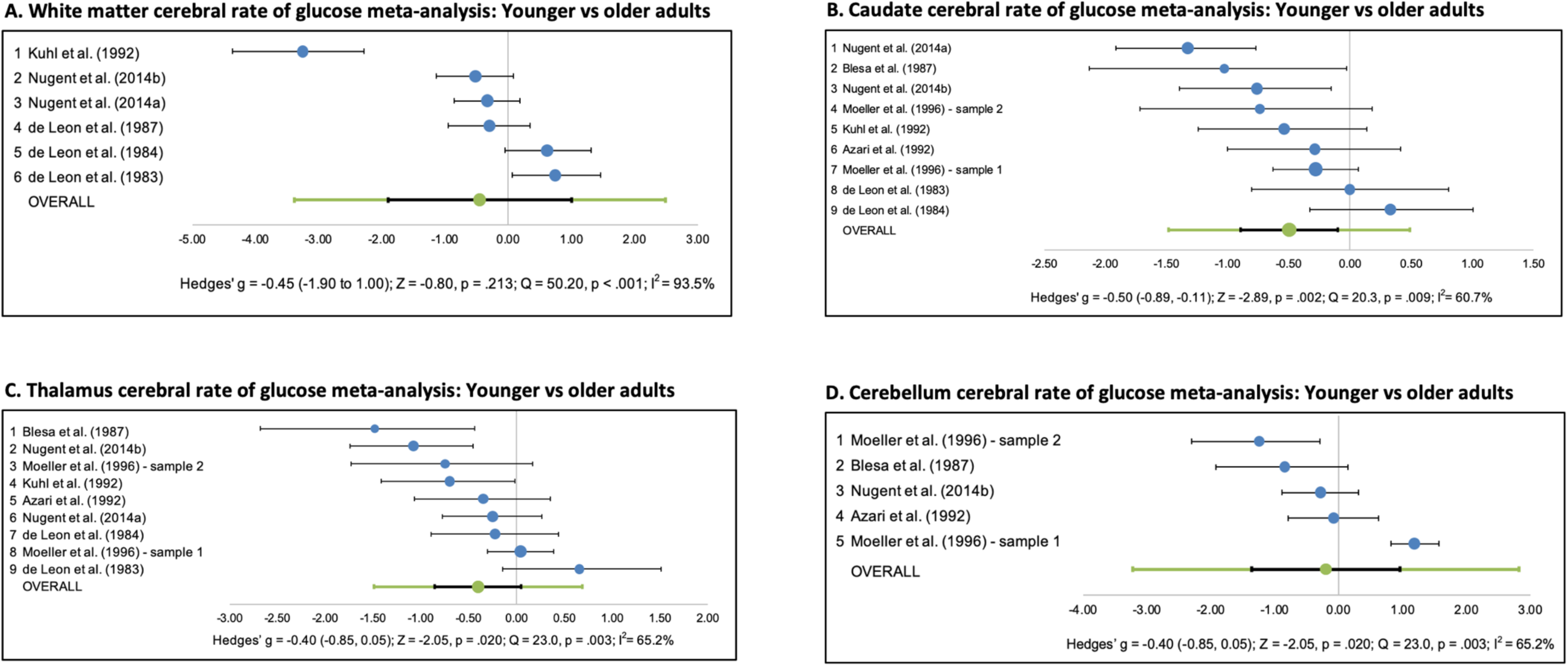
White matter and sub-cortical structures cerebral rate of glucose meta-analyses for younger vs older adults; A. White matter; B. Caudate; C. Thalamus; D. Cerebellum. For each study, the blue dot show Hedges’ g effect size on the X-axis and the size of the dot the weight of the study in the overall effect size. The horizontal line illustrates the high and low limits of the 95% confidence interval. The overall effect size is shown in the bottom row, with Hedges’ g effect size (green dot) and its confidence interval (black) and its prediction interval (green). The prediction interval gives the 95% confidence range in which the outcome of a future study will fall, assuming that the effect sizes are normally distributed.

We performed two separate analyses; the first examined the regional distribution of peak activation where glucose uptake was lower in older than younger adults. The second analyses examined the regional distribution of peak activation where glucose uptake was greater in older than younger adults.

#### 2.5.3. Additional Analyses

We undertook additional analyses on the CRM_GLC_ effect sizes to investigate the potential impact of averaging regions that were not always of the same size. We also undertook contrast analysis in GingerALE to test the PVE-corrected and uncorrected sets of foci for statistically significant differences in spatial convergence.

## Results

### 3.1 Included Studies and Samples

#### 3.1.1 Pooled Effect Size Meta-Analyses

Twenty-two studies with 24 samples (n = 993 participants) met the inclusion criteria and were retained in the systematic review and effect size meta-analyses. The studies were published between 1982 and 2021. The studies are summarised in Table 1. Studies sample sizes ranged from eight to 130 participants. Ten studies with 13 samples reported correlational analysis of age and CMR_GLC_ at a whole-brain level, with the samples covering adults in the second to ninth decade of life. Nineteen studies with 20 samples reported age-group differences. Fifteen of those studies reported age group differences for younger adults with an age range or mean age in the second to third decade of life compared to older adults with an age range or mean age in the sixth to ninth decade of life. One study (Kushner et al., 1987) included adult aged 47-73 yeas in the older age group. Three studies split the samples into younger and older groups with the division at age 50 years (Blesa et al., 1997; Moeller et al., 1996; Yoshi et al., 1998).

The samples included in the two meta-analyses were largely independent. In particular, of the 22 studies in the effect-size meta-analysis, two also reported coordinates of peak metabolic age-group differences and were included in the ALE analysis (Ibanez et al., 2004; Petit-Taboue et al., 1998).

#### 3.1.2 ALE Meta-Analyses

Eight studies with 11 samples (n = 713 participants) published between 1998 and 2014 were included in meta-analysis (see Table 2). The PRISMA process did not identify studies published after 2014 in which the location of peak age group differences in CRM_GLC_ were reported. Studies sample size ranged from 24 to 234 participants. Seven studies included participants from the second or third decades to the eighth or ninth decade of life, and one study included participants from the third to the eighth decade (Yoshizawa et al., 2014).

### 3.2 Cerebral Rate of Glucose Effect Sizes

Table 3 summarises the results of the pooled effect size meta-analyses. Older adults had a 7% lower CRM_GLC_ than younger adults across the whole brain (Hedges’ g = -0.47, Z = -3.73, p < .001). A significant negative correlation was also found between whole-brain CRM_GLC_ and age (Hedges’ g = - 0.24, Z = -5.22, p < .001). The forest plots (Figures 2) show that the studies were moderately heterogenous for age group differences (Q = 25.9, p< .05; l^2^ = 45.9%) but not for the correlation with age (Q = 13.9, p = .305; l^2^ = 13.8%).

**Table 3.** Summary of meta-analyses findings. For Hedges’ g, numbers in parentheses are 95% confidence intervals.

|  | Samples | N-Size | Mean % Reduction<br>Younger vs older | Hedge's G<br>Effect Size | Z-Value | Heterogeneity Q | Heterogeneity (I<sup>2</sup>) |
| --- | --- | --- | --- | --- | --- | --- | --- |
| **Whole brain: Correlation with Age** | 13 | 567 |  | -0.24 (-0.33, -0.14) | -5.22, p = .001 | 13.90, p = .305 | 13.8% |
| **Whole brain: Age Group differences** | 15 | 620 | 7% | -0.47 (-0.74, -0.20) | -3.73, p = .000 | 25.89, p = .027 | 45.9% |
| **Cortex: Age Group Differences** |  |  |  |  |  |  |  |
| Frontal | 13 | 561 | 12% | -0.66 (-0.89, -0.43) | -6.30, p = .000 | 15.06, p = .238 | 20.3% |
| Temporal | 12 | 546 | 11% | -0.56 (-0.90, -0.22) | -3.67, p = .000 | 24.46, p = .011 | 55.0% |
| Parietal | 13 | 559 | 9% | -0.37 (-0.72, -0.02) | -2.30, p = .011 | 29.85, p = .003 | 59.8% |
| Occipital | 8 | 414 | 8% | -0.39 (-0.64, -0.14) | -3.63, p = .000 | 7.60, p = .370 | 7.8% |
| **White Matter: Age Group differences** | 6 | 265 | 3% | -0.45 (-1.90, 1.00) | -0.80, p = .213 | 50.20, p < .001 | 90.4% |
| **Subcortical: Age Group differences** |  |  |  |  |  |  |  |
| Caudate | 9 | 400 | 9% | -0.50 (-0.89, -0.10) | -2.89, p = .002 | 20.34, p = .009 | 60.7% |
| Thalamus | 9 | 400 | 8% | -0.40 (-0.85, 0.05) | -2.05, p = .020 | 23.00, p = .003 | 65.2% |
| Cerebellum | 5 | 243 | 3% | -0.20 (-1.36, 0.96) | -0.48, p = .317 | 42.4, 1 p < .001 | 90.6% |

Significant lower CMR_GLC_ among older adults was found for the frontal, temporal, occipital and parietal lobes. The average reduction in CRM_GLC_ from younger to older adults was 12% for the frontal lobe, (Hedges’ g = -0.66, Z = -6.30, p < 001), 11% for the temporal lobe (Hedges’ g = -0.56, Z = -3.67, p< 001), 9% for the parietal lobe (Hedges’ g = -0.39, Z = -3.63, p < .001) and 8% for the occipital lobe (Hedges’ g = -0.37, Z = -2.30, p < .05). The forest plot for the frontal and occipital lobes (Figures 3A, D) shows low heterogeneity among the studies (frontal, Q = 15.06, p = .238; I^2^ = 20.3%; occipital, Q = 7.60, p = .370, I^2^ = 7.8%). However, for the temporal and parietal lobes (Figures 3B, C), heterogeneity of the studies was moderate and significant (temporal, Q = 24.46, p = .011’ I^2^ = 55.0%; parietal, Q = 29.85, p = .003, I^2^ = 59.8%).

The caudate showed 9% lower glucose metabolism in older adults (Hedges’ g = -0.50, -2.89, p < .01). An 8% reduction was observed in the thalamus, although the confidence interval marginally crossing zero (-0.85 to 0.05) (see Figures 4C). The heterogeneity among the studies was moderate-to-large and significant for each sub-cortical structure (caudate, Q = 20.3, p = .009, I^2^ = 60.7%; thalamus, Q = 23.0, p = 003, I^2^ = 65.2%; cerebellum, Q = 42.4, p < .001, I^2^ = 90.5%). White matter CMR_GLC_ was similar in older and younger adults (Hedges’ g = -0.45 4, Z = -0.80, p = .213).

### 3.3 Cerebral Rate of Glucose Effect Sizes: Volume-Correction vs No Correction

Table 4 summarises the sub-group analyses comparing studies correcting for volume and those with no volume correction. For the volume-correction sub-group, all studies were used that adopted either partial volume effect correction, a measure of atrophy, and/or registration of PET images to MR or CT (see Table 1).

**Table 4:** Sub-group analyses for studies with and without brain volume correction in ageing. Numbers in parentheses are 95% confidence intervals.

|  | Between Sub-Group Heterogeneity ANOVA | Sub-Group Analyses |  |  |  |  |  |  |  |  |  |  |  |
| --- | --- | --- | --- | --- | --- | --- | --- | --- | --- | --- | --- | --- | --- |
|  |  | No correction for volume |  |  |  |  |  | Correction for volume |  |  |  |  |  |
|  |  | Samples | N-size | Mean % Reduction: Younger vs Older | Effect Size | Heterogeneity Q | I<sup>2</sup> | Samples | N-size | Mean % Reduction: Younger vs Older | Effect Size | Heterogeneity Q | I<sup>2</sup> |
| Global | p = .017 | 8 | 355 | 10% | -0.69 (-0.96, -0.42) | 9.49, p = .219 | 26.3% | 7 | 265 | 4% | -0.19 (-0.51, 0.13) | 9.06, p = .170 | 33.8% |
| Cortex |  |  |  |  |  |  |  |  |  |  |  |  |  |
| Frontal | p = .303 | 6 | 309 | 13% | -0.76 (-1.07, -0.46) | 7.12, p = .212 | 29.8% | 7 | 252 | 11% | -0.55 (-0.83, -0.27) | 6.86, p =.334 | 12.5% |
| Temporal | p = .042 | 6 | 310 | 14% | -0.79 (-1.08, -0.49) | 7.35, p = .196 | 32.0% | 6 | 236 | 6% | -0.27 (-0.70, 0.15) | 10.07, p = .073 | 50.4% |
| Parietal | p = .025 | 6 | 165 | 12% | -0.62 (-0.82, -0.41) | 3.95, p = .557 | 0.0% | 7 | 120 | 5% | -0.34 (-0.78, 0.10) | 10.38, p =.055 | 53.8% |
| Occipital | p = .898 | 5 | 293 | 9% | -0.44 (-0.57, -0.30) | 1.30, p =.862 | 0.0% | 3 | 121 | 7% | -0.39 (-1.17, 0.38) | 5.61, p =.061 | 64.4% |
| Subcortical |  |  |  |  |  |  |  |  |  |  |  |  |  |
| Caudate | p = .594 | 4 | 217 | 8% | -0.36 (-0.53, -0.19) | 1.22, p = .749 | 0.0% | 5 | 183 | 10% | -0.55 (-1.18, 0.07) | 17.84, p = .001 | 77.6% |
| Thalamus | p = .785 | 4 | 217 | 6% | -0.33 (-0.72, 0.05) | 5.68, p =.128 | 47.2% | 5 | 183 | 10% | -0.44 (-1.14, 0.26) | 16.33, p =.003 | 75.5% |
| Cerebellum | p = .551 | 3 | 182 | -1% | 0.01 (-1.37, 1.39) | 27.95 p = .000 | 92.8% | 2 | 61 | 12% | -0.44 (-0.94, 0.06) | 0.97, p =.324 | 0.0% |

Significant sub-group effects were found at the whole brain level (p = 0.017), with the effect size larger for the whole brain in the uncorrected (-.69) versus the corrected sub-groups (-.19) and corresponding to a 10% and 4% average reduction, respectively (see Figure 5a). Heterogeneity was relatively low for both sub-groups (uncorrected, l^2^ = 33.8%; corrected, l^2^ = 26.3%). Significant sub-group effects were also found or the temporal (p = .042) and parietal (p = .025) lobes, with the effect sizes significant for uncorrected but not corrected samples (confidence intervals crossing zero). The reductions for the corrected samples were 6% in the temporal lobe, 5% in the parietal lobe and 7% in the occipital lobe.

**Figure 5.**
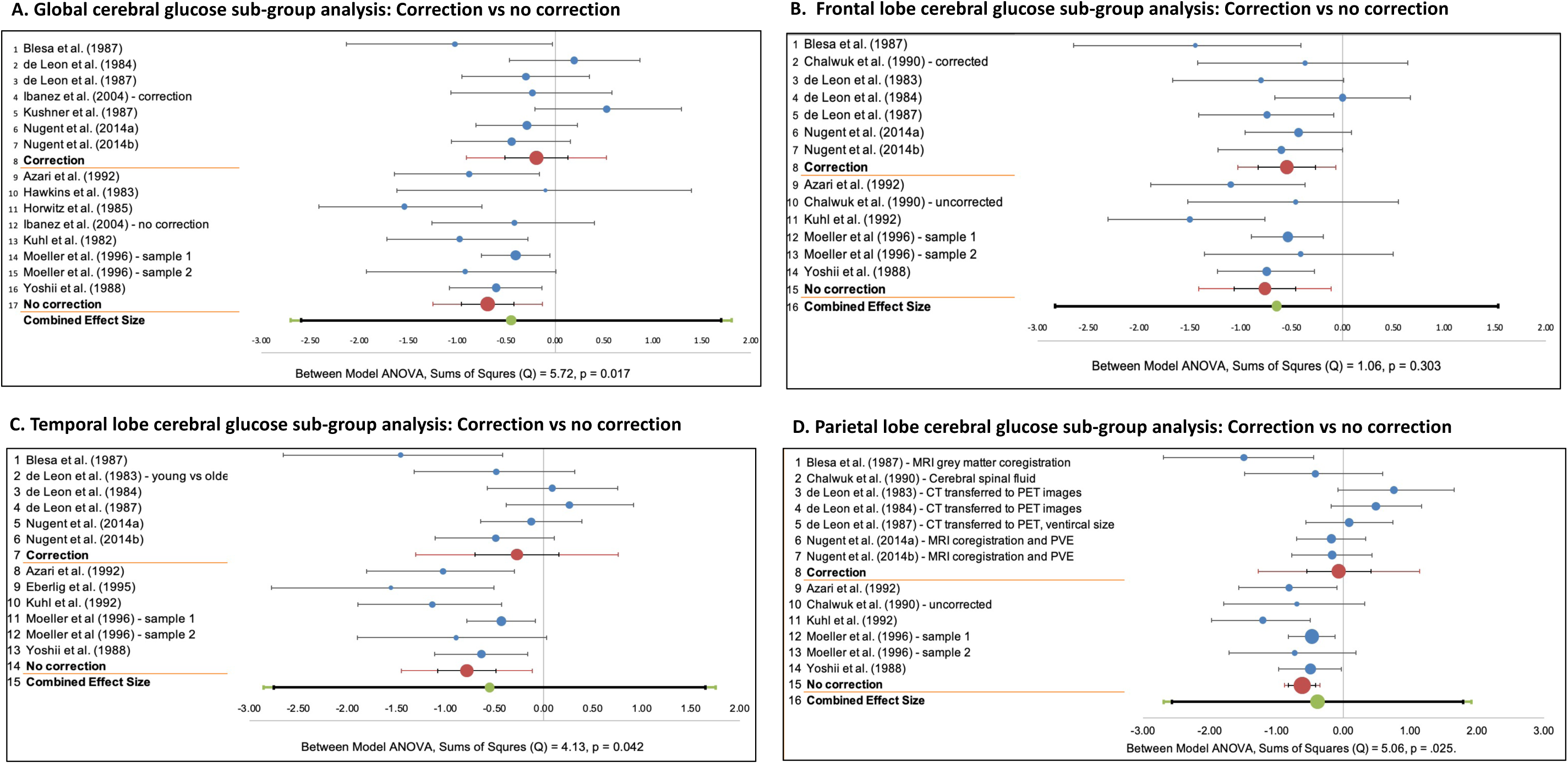
Sub-group analyses for brain areas showing significant effect size differences for volume correction vs no correction. A. Whole brain; B. Frontal lobe; C. Temporal lobe; D. Parietal lobe. Blue dots represent individual study effect sizes on the X-axis and their weight (size of dot) in the pooled effect; red dots represent subgroups effect sizes, and the green dot represents the combined effect size. For each study, the horizontal line shows the high and low limits of the 95% confidence interval. The confidence interval is shown in black for the subgroups and combined effect size and the prediction intervals are shown in their respective colours.

Forest plots for the sub-group analyses that showed significant differences for corrected versus uncorrected effect sizes, as well as the frontal lobe, are shown in Figure 5. For the frontal lobe, no significant sub-group difference was found (p = .303). In other words, even after correcting for volume, partial volume effects, or both, older adults had a lower cerebral rate of glucose in the frontal lobe that is not statistically different in effect size to uncorrected rates (see Figure 5B). The average lower CRM_CLC_ among older adults was 13% in the uncorrected samples and 11% in the corrected samples.

For the occipital lobe, no significant sub-group difference was found (p = .898). Only the uncorrected effect size was significant (95% CI -0.57, -0.30). Heterogeneity was higher among the corrected than uncorrected samples for the temporal (uncorrected, l^2^ = 32.0%; corrected l^2^ = 50.4%), the parietal (uncorrected, l^2^ = 0%; corrected l^2^ = 53.8%) and the occipital (uncorrected, l^2^ = 0%; corrected l^2^ = 64.4%) lobes.

Sub-group differences were not evident in the sub-cortical structures (caudate, p = .594; thalamus, p = .785; cerebellum, p = .551), although the caudate confidence interval was significant for the uncorrected studies (95% CI, -0.53 to -0.19) and marginally spanned zero for the volume-corrected studies (95% CI, -1.18 to 0.07). Heterogeneity was higher among the corrected than uncorrected samples for the caudate (uncorrected, l^2^ = 0%; corrected l^2^ = 77.6%), the thalamus (uncorrected, l^2^ = 47.2%; corrected l^2^ = 75.5%), but not the cerebellum (uncorrected, l^2^ = 92.8%; corrected l^2^ = 0%). The reduction in CRM_GLC_ for the caudate, thalamus and cerebellum was higher in the corrected samples than the uncorrected samples (10% versus 8% for the caudate; 10% versus 6% for the thalamus and 12% versus -1% for the cerebellum).

Insufficient data was available to undertake sub-group analyses for white matter and the whole-brain correlation with age.

### 3.4 ALE: Areas of Lower Metabolism in Older than Younger Adults

All eight studies and 11 samples reported lower peak activations in older than younger adults. The ALE analysis identified nine clusters of lower metabolism in older adults ranging in size from 200mm^3^ to 2,640mm^3^ (see Table 5; Figures 6-9). The largest clusters (clusters 1 and 2) are mostly in the left and right inferior frontal and superior temporal gyri and the insula. A third, relatively large cluster is mostly in the inferior frontal gyrus (cluster 4), with activity also in the middle frontal and precentral gyri. Relatively large clusters are also found in the anterior cingulate and cingulate gyrus (clusters 3 and 8). Smaller clusters are in the caudate (cluster 5), mostly the subcallosal gyrus (cluster 6), the inferior parietal and superior temporal gyri (cluster 7) and thalamus (cluster 9).

**Figure 6.**
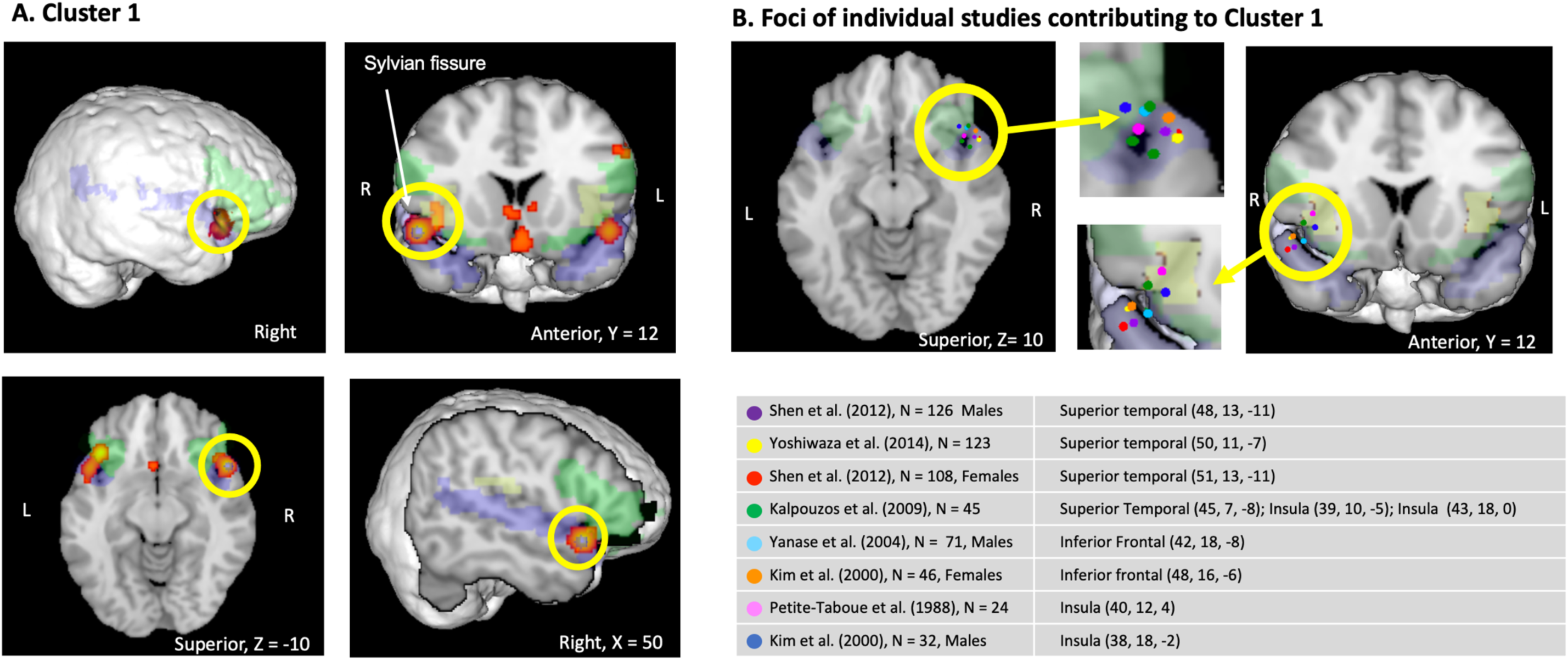
Clusters of lower glucose metabolism among older versus younger adults from the ALE meta-analysis. A. Right, anterior and superior views, with 3mm diameter blue ROI placed at maximum ALE value is in the superior temporal gyrus (50, 12, -10). Superior temporal gyrus shaded blue; inferior frontal gyrus shaded green and Insula shaded yellow. The lateral fissure (Sylvian fissure) is shown separating the frontal and temporal gyri. B. Plot of individual study foci for Cluster 1, showing activity in the superior temporal and inferior frontal gyri and the insula.

**Table 5.**
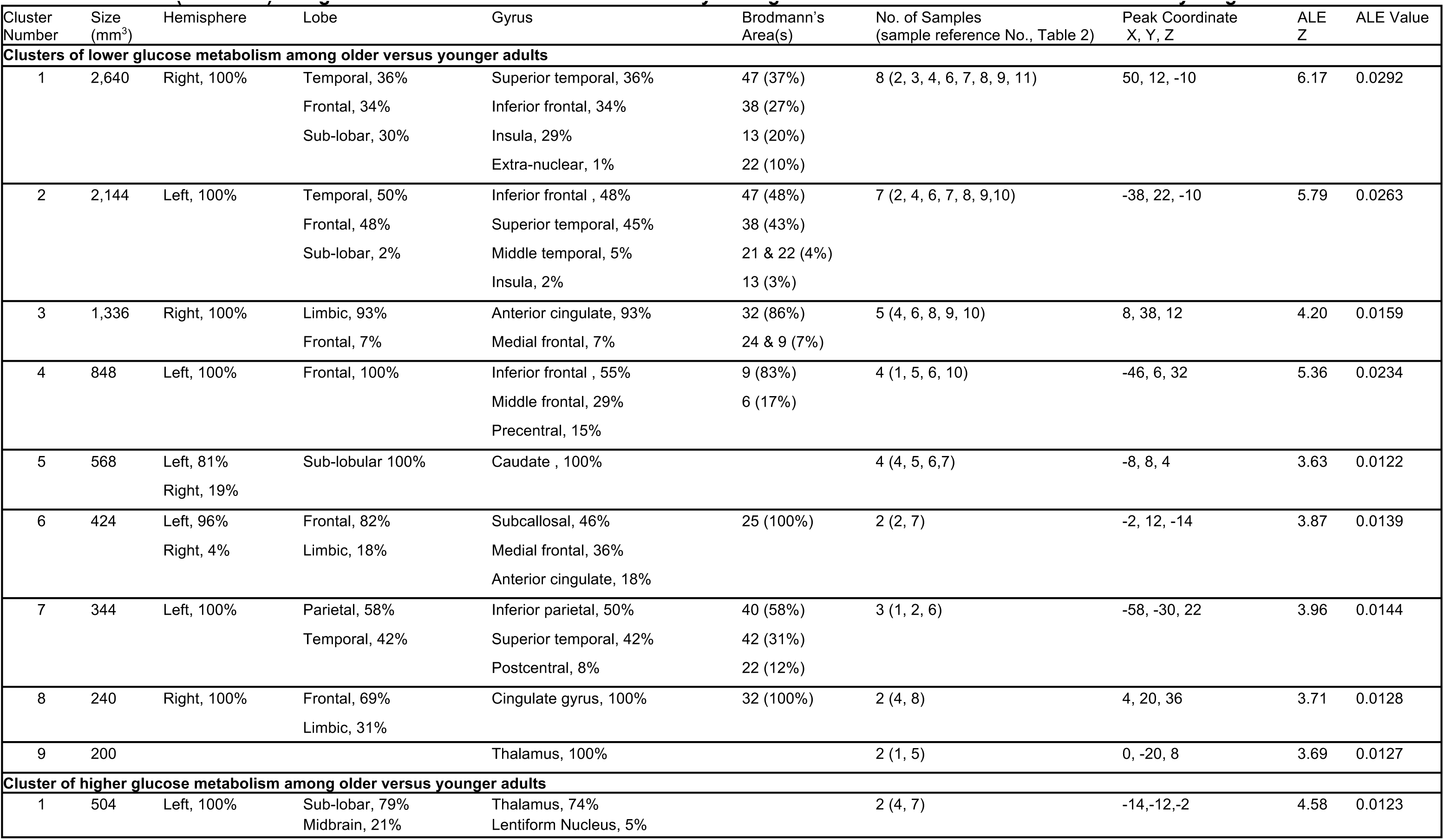
Locations (Talairach) of significant clusters from the ALE meta-analysis of glucose metabolism differences for older vs younger adults

| Cluster Number | Size (mm<sup>3</sup>) | Hemisphere | Lobe | Gyrus | Brodmann's Area(s) | No. of Samples (sample reference No., Table 2) | Peak Coordinate X, Y, Z | ALE Z | ALE Value |
| --- | --- | --- | --- | --- | --- | --- | --- | --- | --- |
| **Clusters of lower glucose metabolism among older versus younger adults** |  |  |  |  |  |  |  |  |  |
| 1 | 2,640 | Right, 100% | Temporal, 36%<br>Frontal, 34%<br>Sub-lobar, 30% | Superior temporal, 36%<br>Inferior frontal, 34%<br>Insula, 29%<br>Extra-nuclear, 1% | 47 (37%)<br>38 (27%)<br>13 (20%)<br>22 (10%) | 8 (2, 3, 4, 6, 7, 8, 9, 11) | 50, 12, -10 | 6.17 | 0.0292 |
| 2 | 2,144 | Left, 100% | Temporal, 50%<br>Frontal, 48%<br>Sub-lobar, 2% | Inferior frontal, 48%<br>Superior temporal, 45%<br>Middle temporal, 5%<br>Insula, 2% | 47 (48%)<br>38 (43%)<br>21 & 22 (4%)<br>13 (3%) | 7 (2, 4, 6, 7, 8, 9, 10) | -38, 22, -10 | 5.79 | 0.0263 |
| 3 | 1,336 | Right, 100% | Limbic, 93%<br>Frontal, 7% | Anterior cingulate, 93%<br>Medial frontal, 7% | 32 (86%)<br>24 & 9 (7%) | 5 (4, 6, 8, 9, 10) | 8, 38, 12 | 4.20 | 0.0159 |
| 4 | 848 | Left, 100% | Frontal, 100% | Inferior frontal, 55%<br>Middle frontal, 29%<br>Precentral, 15% | 9 (83%)<br>6 (17%) | 4 (1, 5, 6, 10) | -46, 6, 32 | 5.36 | 0.0234 |
| 5 | 568 | Left, 81%<br>Right, 19% | Sub-lobular 100% | Caudate, 100% |  | 4 (4, 5, 6, 7) | -8, 8, 4 | 3.63 | 0.0122 |
| 6 | 424 | Left, 96%<br>Right, 4% | Frontal, 82%<br>Limbic, 18% | Subcallosal, 46%<br>Medial frontal, 36%<br>Anterior cingulate, 18% | 25 (100%) | 2 (2, 7) | -2, 12, -14 | 3.87 | 0.0139 |
| 7 | 344 | Left, 100% | Parietal, 58%<br>Temporal, 42% | Inferior parietal, 50%<br>Superior temporal, 42%<br>Postcentral, 8% | 40 (58%)<br>42 (31%)<br>22 (12%) | 3 (1, 2, 6) | -58, -30, 22 | 3.96 | 0.0144 |
| 8 | 240 | Right, 100% | Frontal, 69%<br>Limbic, 31% | Cingulate gyrus, 100% | 32 (100%) | 2 (4, 8) | 4, 20, 36 | 3.71 | 0.0128 |
| 9 | 200 |  |  | Thalamus, 100% |  | 2 (1, 5) | 0, -20, 8 | 3.69 | 0.0127 |
| **Cluster of higher glucose metabolism among older versus younger adults** |  |  |  |  |  |  |  |  |  |
| 1 | 504 | Left, 100% | Sub-lobar, 79%<br>Midbrain, 21% | Thalamus, 74%<br>Lentiform Nucleus, 5% |  | 2 (4, 7) | -14, -12, -2 | 4.58 | 0.0123 |

Cluster 1 is in the right hemisphere, with the maximum ALE value in the superior temporal gyrus (50, 12, -10). The cluster includes activation in the superior temporal gyrus (36%), the inferior frontal gyrus (34%) and the insula (29%) (see Figure 6A). The cluster crosses the Sylvian fissure, suggesting that the cluster at least partly reflects the ALE method of placing a kernel at the activity close in space. We further investigated the composition of the cluster by plotting the location of the activation from each contributing study (see Figure 6B). The three largest samples reported lower metabolism among older adults in the superior temporal gyrus, in line with the cluster peak. There are also studies with coordinates in the inferior frontal gyrus and insula. This confirms that the cluster is reflecting lower metabolic activity among older adults in the three distinct regions that are separated in space by the Sylvian Fissure.

Cluster 2 is in the left hemisphere, almost contralateral to cluster 1 (see Figure 7A). It is slightly more anterior and superior to cluster 1, with a maximum ALE value in the inferior frontal gyrus (-38, 22, -10). The cluster includes areas of the inferior frontal gyrus (45%), superior temporal gyrus (45%), medial temporal gyrus (15%) and insula (2%). Three of the larger studies reported lower metabolism among older adults in the inferior frontal gyrus, whereas one large study reported lower metabolism in the superior temporal gyrus (see Figure 7B). Cluster 4 also has a maximum ALE value and highest percentage of activity (55%) in the left inferior frontal gyrus (-46, 6, 32). It also includes activity in the middle frontal (29%) and precentral (15%) gyri (see Figure 4).

**Figure 7.**
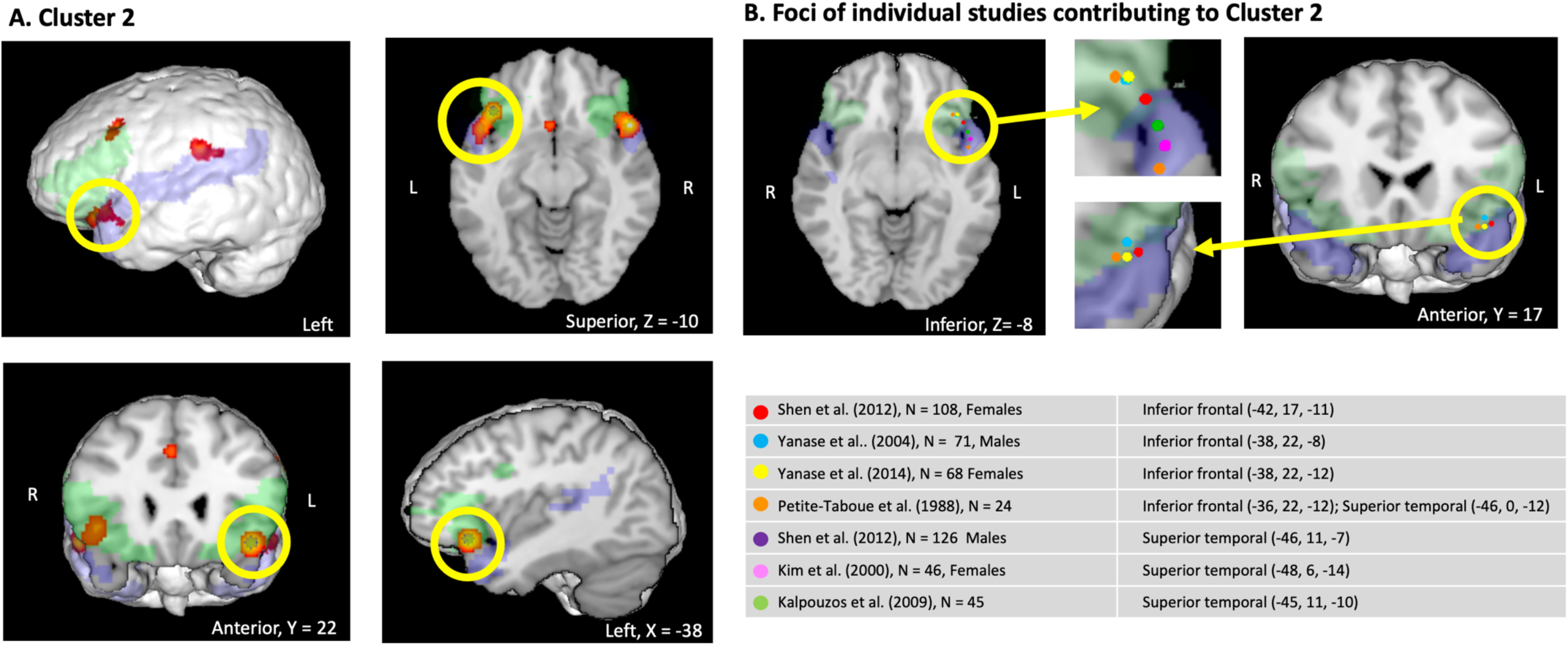
Clusters of **lower** glucose metabolism among older versus younger adults from the ALE meta-analysis. A. Left, superior and anterior views, with 3mm diameter blue ROI placed at maximum ALE value is in the inferior frontal gyrus (-38, 22, -10). Superior temporal gyrus shaded blue and inferior frontal gyrus shaded green. B. Plot of individual study foci for Cluster 2, showing activity in the inferior frontal and superior temporal gyri.

Cluster 3 is predominantly in the anterior cingulate (93%), with a smaller amount of activity in the medial frontal gyrus (7%) (see Figure 8). Cluster 8 is located (100%) in the cingulate gyrus, with a peak ALE value at (4, 20, 36). Cluster 5 is located (100%) in the caudate (see Figure 9). Cluster 6 has its peak location (-2, 12, -14) and largest percentage of activation (46%) in the subcallosal gyrus, with activity also in the medial frontal (36%) and anterior cingulate (18%). Cluster 7 has its peak (-58, -30, 22) and highest percentage (50%) of activity in left inferior parietal lobule, with 42% of activity in the superior temporal gyrus and 8% in the postcentral gyrus (see Figure 9). Cluster 9 is (100%) in the thalamus (Figure 9).

**Figure 8.**
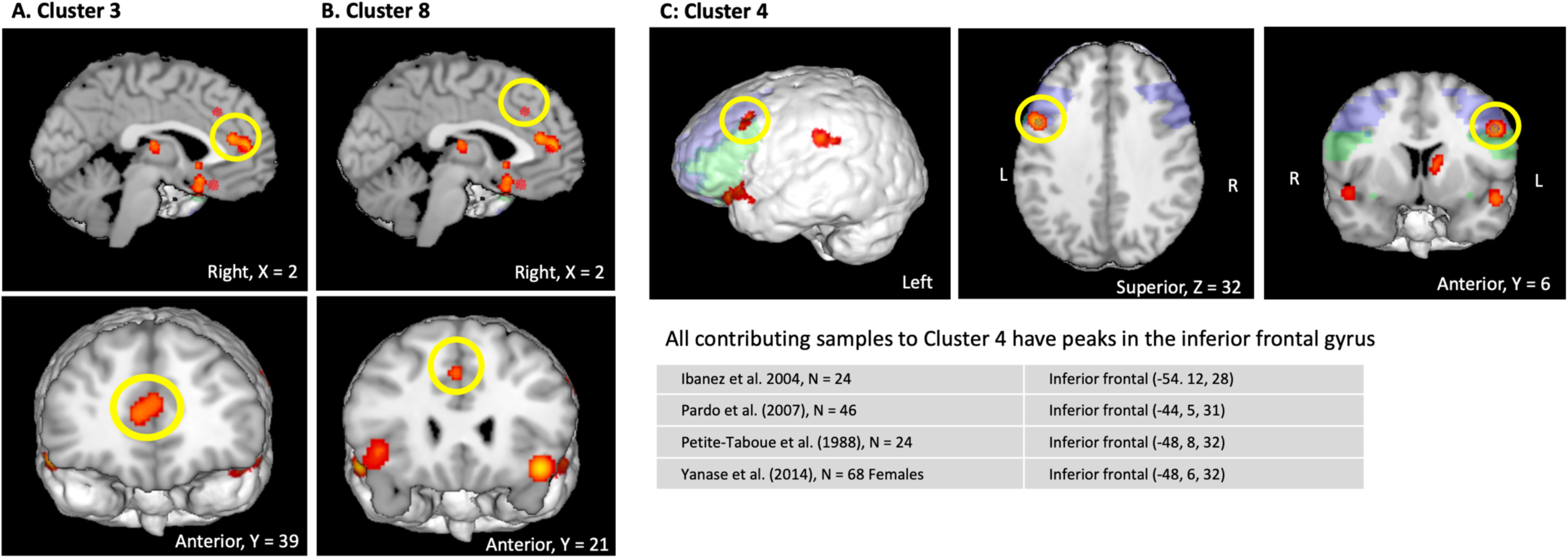
Clusters of **lower** glucose metabolism among older versus younger adults from the ALE meta-analysis. A. Cluster 3 in the anterior cingulate. B. Cluster 8 in the cingulate gyrus. C. Cluster 4, with 3mm diameter blue ROI placed at maximum ALE value is in the inferior frontal gyrus (-46, 6, 32). Activity is 55% inferior frontal gyrus (green shading), 29% middle frontal gyrus (blue sharing) and 15% precentral gyrus. Individual studies showing peak activity in the inferior frontal gyrus. For cluster 4, contributing studies are shown to illustrate all peaks are in the inferior frontal gyri. For studies contributing to clusters 3 and 9, see Tables 5, column headed “Samples No.”, and the corresponding study numbers in Table 2.

### 3.5 Areas of Higher Metabolism in Older than Younger Adults

Three studies (Kim et al., 2009; Shen et al., 2012; Yanase et al., 2005) with five samples reported locations where metabolism was higher for older versus young adults. The ALE analysis identified one cluster in which glucose metabolism is higher for older versus younger adults (see Table 5 and Figure 9E). The cluster is 504mm^3^ and is in the left hemisphere in the thalamus (74%) and lentiform nucleus (5%).

**Figure 9.**
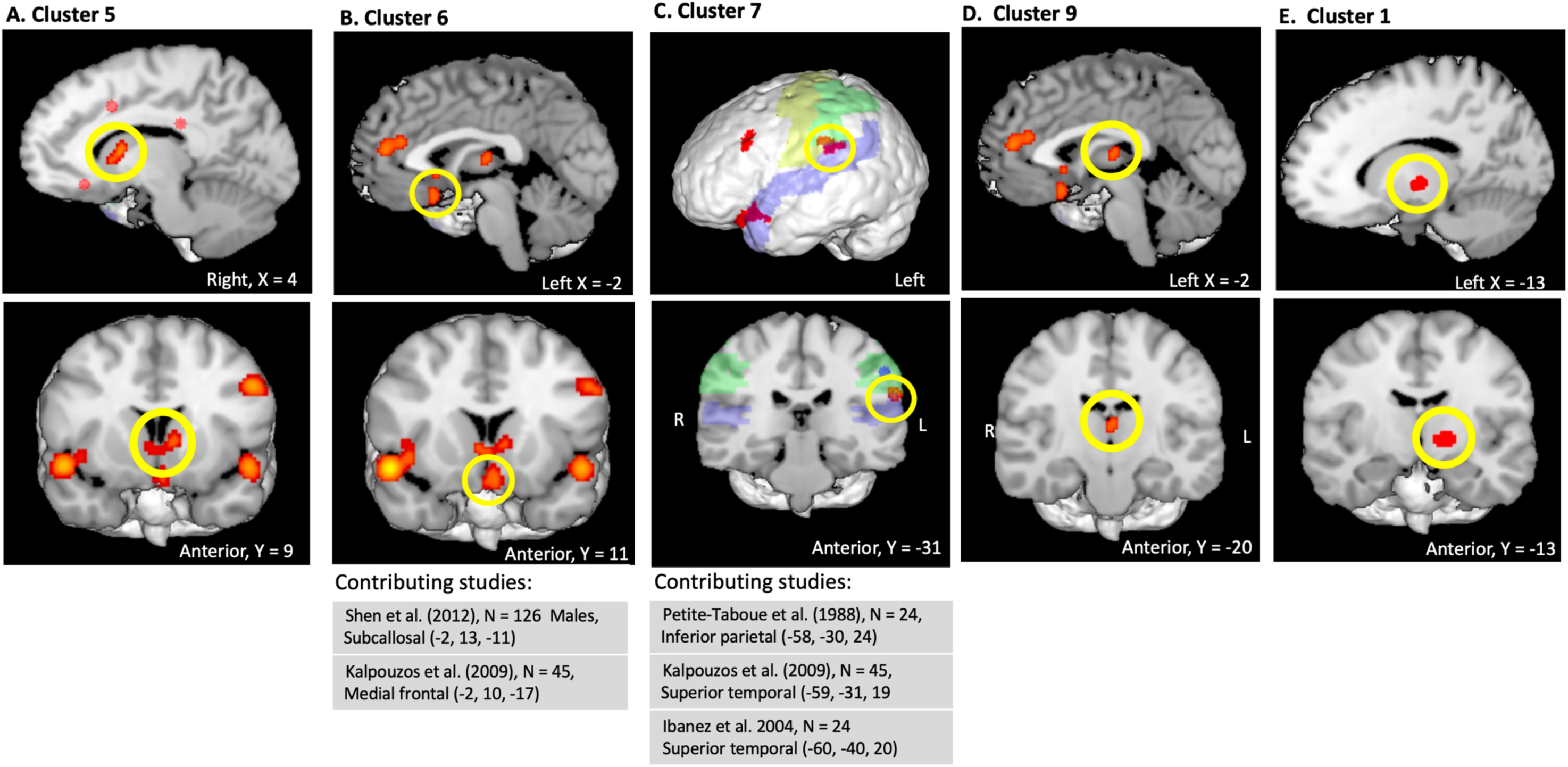
Clusters of age differences in glucose metabolism among older versus younger adults from the ALE meta-analysis. Lower glucose metabolism: A. Cluster 5 in the caudate. B. Cluster 6, 46% subcallosal gyrus, 36% medial frontal gyrus, 18% anterior cingulate. C. Cluster 7, 50% inferior parietal lobule (green shading), 42% superior temporal gyrus (blue shading), 8% postcentral gyrus (yellow shading). D. Cluster 9 in the thalamus. Higher glucose metabolism: E. Cluster 1, 74% Thalamus, 5% Lentiform Nucleus. For cluster 6 and 7, contributing studies are shown to illustrate that peaks span multiple gyri. For studies contributing to clusters 5, 9 and 1, see Tables 5, column headed “Samples No.”, and the corresponding study numbers in Table

### 3.6 Additional Analyses

The analyses to investigate the impact of averaging ROIs yielded results consistent with those reported in section 3.1 and 3.2. Results are shown in the Supplement. Specifically, the strongest age-related metabolic reduction was found in the frontal lobe and it remained significant after volume correction. Moreover, the corrected vs uncorrected results across the other lobes are consistent with the results reported for all samples (see Tables S1 and S2).

Results for the contrast analyses comparing the PVE-corrected and uncorrected samples showed three areas of conjunction in the inferior and medial frontal gyri, overlapping clusters 1, 2 and 4 from the analysis of the combined samples (see Table S3 and S4 and Figure S1A-C).

## 4. Discussion

### 4.1 Glucose Hypometabolism in Ageing is Predominantly Frontal and Temporal

Using two separate but complementary analytical techniques in two largely independent samples, we found older people tended to have lower frontal and temporal glucose metabolism than younger people. This pattern existed when the analysis was performed at the lobular levels as well as at the more granular level using ALE whereby older people had lower glucose metabolism in the inferior frontal, superior temporal and inferior temporal junction bilaterally. Taken together the meta-analyses suggest that ageing from the third to seventh decade of life is associated with glucose hypometabolism that is predominantly frontal and temporal. A general hypometabolism at the lobular level is accompanied by specific and significant regional changes in the frontal and temporal lobes. The effect-size meta-analysis of over 600 participants found that the negative effect of age on glucose metabolism was greatest in the frontal lobe, corresponding to a 11% reduction after volume correction. A 6% reduction was found in the temporal lobe. In the ALE meta-analysis of over 700 participants, we found that older adults had lower cerebral glucose metabolism than younger adults in areas of the left and right inferior frontal and superior temporal gyri and the inferior temporal junction.

The results of the meta-analyses are consistent with literature indicating that the ageing brain displays impaired glucose uptake and metabolism. The reasons why the older brain may show reduced glucose uptake are unclear but may include an impairment in neurons involved in glucose transport, the loss of glial energetics to support neurons, the attenuation of neurovascular coupling, and loss of enzyme function involved in energy production via the Krebs cycles and aerobic glycolysis (for reviews see Cunnane et al., 2020; Kapogiannis & Avgerinos, 2020). Cerebral metabolism is also influenced by the endocrine modulation of appetite and whole-body energy supply, processes that are also compromised during normal ageing and dementia (Cunnane et al., 2020). Brain insulin is also implicated as a mediator of altered glucose metabolism in ageing through several means, including altering glucose transport. This may be due to poor signalling of insulin receptors, reduced insulin levels in the brain, or reduced transportation of insulin into the brain. Evidence of decreased brain insulin receptor number and function has been reported in both clinical samples and animal models of aging and dementia (Frazier et al., 2019).

### 4.2 Glucose Hypometabolism in Ageing is Mediated by Brain Volume Loss

Our results indicate that correcting for brain volume changes and partial volume effects in PET imaging influences the sensitivity and effect size estimation of age-related CMR_GLC_ differences. Previous research indicates that that prefrontal cortex and temporal lobes are the most affected by atrophy in ageing (Zanto & Gazzley, 2019); whereas the occipital lobes show the least volume changes (Toepper, 2017). In the current effect size meta-analyses, the corrected samples showed substantially lower effect sizes than the uncorrected samples for the temporal lobe (14% uncorrected; 6% corrected) but less so in the occipital lobe (9% uncorrected; 7% corrected). These results suggest that glucose metabolism is partly although not entirely related to changes in brain volume in ageing, particularly in the lobes most vulnerable to volume loss.

Contrast analysis provided additional support for the strength of the hypometabolism in ageing in the left and right inferior frontal and left superior temporal gyri (see supplementary materials). These locations of hypometabolism were evident in the PVE-corrected samples. In the case of the frontal regions, they were also evident in the uncorrected samples. Statistically separate but spatially similar regions of hypometabolism in ageing were also found in the anterior cingulate in the PVE-corrected and uncorrected samples, suggesting that the anterior cingulate is also vulnerable to metabolic reductions in ageing.

A cluster in the non PVE-corrected samples only was found in the caudate, and a second cluster in the cingulate. It is possible that the presence of clusters in the uncorrected samples only is at least partly an artifact of measurement error from lack of PVE correction. It is also possible that PVE or volume correction will have less of an impact on the measured rates of glucose in the sub-cortical structures. The pooled effect sizes were larger in the corrected than uncorrected samples for the caudate, thalamus and cerebellum (Table 4). The larger reduction in the corrected samples may reflect several factors. First, these structures tend to show relatively low volume and metabolic changes with age compared to many cortical areas (e.g., Kawasaki et al., 2008; Viglianti et al. (2019). Second, we also found large heterogeneity among the corrected samples for the caudate and the thalamus in the pooled effect size analyses. Taken together, these results suggest that there is significant variability in the age-related metabolic effects in the sub-cortical regions in the available studies, and that additional research is needed to assess the effect sizes and locations of metabolic changes in older adults in those structures.

### 4.3 Age-related hypometabolism in sub-cortical structures

The effect size meta-analysis showed lower glucose metabolism among older adults in the caudate (10% after volume correction; CI, -1.18 to 0.07). In the ALE meta-analysis, we also found that older adults had lower metabolism in regions of the caudate, although only in the non PVE-corrected samples. The caudate is central to the cortical-basal ganglia circuits that are important for many cognitive functions (see Jamadar 2018 for a review), including learning. Connections to the caudate from the prefrontal and anterior cingulate cortices are also crucial for working memory (Grazioplene et al., 2015). Age-related decline in anterior cingulate glucose metabolism (clusters 3 and 8) also correlates with a decline in cognition, including attentional and inhibitory control and verbal fluency (Pardo et al., 2007; Petersen & Posner, 2012). Our results raise the possibility that reduced glucose metabolism in the caudate is contributing to these age-related functional changes in learning and memory.

Our meta-analysis results indicate that alterations to glucose metabolism in the caudate (cluster 5) and thalamus (cluster 9) has the potential for widespread impact on both the functional activity of the structures themselves, as well as their extensive connections to other regions, likely impacting the executive, sensory and emotional functions, as well as the planning and execution of movement (Grahn et al., 2008; Grazioplene et al., 2015; Haber, 2016; Lanciego et al., 2012; see Jamadar, 2019, for a review). Hypometabolism in the thalamus also has the potential to alter its role as a hub for relaying sensory information between different subcortical areas and the cerebral cortex (Halassa & Sherman, 2019; Herrero et al., 2002).

Our results suggest that alterations to brain glucose metabolism in ageing are not adding or contributing to the constellation of the known age-related changes in white matter integrity and function. Age-related changes to white matter can include volume loss, lesions and cortical disconnection (Davis et al., 2009; see Liu et al., 2017 for review). One reason for a lack of age effect may be that CRM_CGL_ in white matter is approximately 25% of the values in grey matter (Berti et al., 2014) and that lower absolute rates of glucose metabolism are less vulnerable to age related decline. The present analysis was limited by the availability of six published studies with too few samples having PVE for sub-group analysis. Further research is needed to quantify age effects in white matter glucose metabolism.

### 4.4 The Pattern of Glucose Hypometabolism in Ageing is Different to that in Dementia

The largest ALE clusters were mostly in the superior temporal and inferior frontal gyri, as well as the inferior frontal junction. These locations differ to those reported in an ALE meta-analysis by He et al. (2015), comparing FDG-PET results in Alzheimer’s and MCI to healthy controls. In particular, the location of the hypometabolism is qualitatively different, being predominately anterior in normal ageing in our results and predominantly posterior in AD and MCI in He et al.’s results.

We found the largest region (cluster 1) spanning the right superior temporal and inferior frontal gyri and insula (peak at 50, 12, -10). He et al. (2015) also found a large region of hypometabolism in the right hemisphere although it was substantially more posterior and superior in the angular, parietal, middle/superior temporal and supramarginal gyri (44, -64, 34). He et al. (2015) found a second right-side region of hypometabolism, although it was also substantially more posterior than those we found in normal ageing, being located in the right inferior/middle temporal and fusiform gyri (54, −40, −22). They also found moderate hypometabolism in a cluster in right middle and superior frontal gyrus (32, 14, 54), which was absent in the results of our analyses.

In terms of the left hemisphere, we found a relatively large region (cluster 2) mostly in the inferior frontal and superior temporal gyri (-38, 22, -10). Although two clusters of hypometabolism were also found in the left hemisphere in patients with AD, the clusters were substantially more posterior. In fact, one was both more posterior and inferior in the inferior/middle temporal, fusiform and parahippocampal gyri (-54, -32, -22); the other was both more posterior and superior in the angular, middle temporal, supramarginal, parietal and superior occipital gyri (-44, -66, 34).

We found a relatively small cluster of hypometabolism in normal ageing located 100% in the anterior portion of the right cingulate gyrus (4, 20, 36). Although hypometabolism was also found in a portion of the cingulate gyrus in patients with AD, the cluster was bilateral and larger (1,331 voxels), with the peak substantially more posterior in the precuneus (2, -42, 30).

In terms of hypometabolism that showed similar spatial locations in our normal ageing analyses and He et al.’s AD cohort, the closest was in the thalamus (0, -20, 8 in our results vs 6, −24, 12 in AD). AD patients also showed a moderately decreased metabolism in left middle frontal and precentral gyri (−28, 18, 40), which was more superior to the cluster 2 in our results (-38, 22, -10).

In MCI, a bilateral cingulate gyrus and left precuneus cluster was also found (6, −58, 24), which like the AD cohort was substantially more posterior to the cluster we found in the cingulate in normal ageing (4, 20, 36). He et al. (2015) also found two relatively small clusters of 66 and 81 voxels in the left hemisphere, one being significantly more posterior to any of our clusters (−42, −62, 18), the other being more superior to cluster 2 in our results (−28, 20, 40). A similar profile was found in the left hemisphere, with a small cluster of 23 voxels in the inferior parietal lobule, and superior temporal and angular gyri (40, −52, 38), substantially more posterior to any of the left side clusters we found in normal ageing; and a small cluster of 30 voxels more superior to our clusters in the right middle frontal gyrus (46, 20, 44).

The temporal pole has been implicated in other neurological diseases, including frontotemporal dementia, temporal lobe epilepsy and schizophrenia (see Ding et al., 2009). Imaging studies of patients with frontotemporal dementia have focussed almost entirely on brain structural changes and have shown a pattern of atrophy and hypoperfusion in the frontal, temporal and fronto-insular regions (Bang et al., 2015), locations that are shared in clusters 1 and 2 of the ALE analysis. Bejanin et al. (2020) reported that patients with frontotemporal dementia show an increase in bilateral temporal, dorsolateral, and medial prefrontal hypometabolism over a 15 months follow-up compare to controls, suggesting that cerebral metabolism is qualitatively and quantitatively different in “normal” older adults and individuals with frontotemporal dementia. Given the limited research on glucose metabolism in frontoparietal dementia, additional studies are needed to distinguish it’s metabolic alterations from “normal” ageing.

### 4.5 Glucose Hypometabolism and Theories of Ageing

The pattern of greatest metabolic decline in the frontal lobe overall and in cluster 1 and 2 specifically is consistent with the retrogenesis or the ‘last-in-first-out’ principle (Davis et al., 2009; Reisberg et al., 2002), which states that the last brain regions to develop are the first to degenerate. It is also consistent with evidence that the early developing regions (e.g., brainstem, cerebellum, motor cortex) are the regions that are largely spared from age-related atrophy (Fjell & Walhovd, 2010; Shaw et al., 2008) and metabolic decline (Fjell et al., 2014). In contrast, the frontal regions become metabolically active during late stages of development and are the most consistently affected by age-related changes (see Fjell et al., 2014 for review). The largest metabolic reduction in ageing in the frontal lobe in our pooled effect size meta-analyses support the ‘last-in-first-out” principle.

The results of the meta-analyses provide partial support for the *frontal ageing hypothesis.* The *frontal ageing hypothesis* (Jackson, 1958; also see Greenwood, 2000) predicts that brain change in ageing selectively effects frontal regions. It further predicts that functions largely dependent on frontal regions (e.g., working memory, decision-making) will decline earlier or faster in ageing, whereas functions largely independent of frontal lobes remain relatively spared. The results of the pooled effect size meta-analyses suggests that hypometabolism is greatest in the frontal regions at the global level, providing support for the notion that the frontal lobe is particularly vulnerable to age-related physiological changes. Although the frontal lobe exhibited the largest age-related hypometabolism, non-frontal lobes in the pooled effect size meta-analysis and regions in the ALE meta-analyses also exhibited declines. To further test whether or not metabolic changes in ageing are consistent with the frontal ageing hypothesis, additional research is needed that directly links metabolic changes to functional changes, particularly those that are largely dependent on the frontal lobe.

The *compensation hypothesis* in ageing postulates that older adults recruit higher levels of neuronal activity in comparison to young subjects in some brain areas to compensate for functional deficits located in other regions (Park et al., 2004; Reuter-Lorenz &, Cappell, 2008; for review, see Ebaid & Crewther, 2020). For example, greater recruitment of prefrontal cortical regions involved in executive functions is also frequently reported in older adulthood compared to younger adults. These frontal recruitment patterns have been described as posterior-to-anterior shift in aging (PASA) (see Cabeza & Dennis, 2012; Davis et al., 2008; Spreng & Turner, 2019, for review of these theories). In contrast to the compensatory hypothesis, the *neural efficiency hypothesis* (Haier et al., 1988) proposes that more efficient brain functioning is evident in cognitively high performing individuals. The more efficient brain function is also associated with lower brain activation compared to lower performing individuals undertaking the same task.

Much of the evidence for the compensation and neural efficiency hypotheses come from functional magnetic resonance imaging (fMRI) studies. However, it is worth noting that the neural efficiency phenomenon was first reported in a PET study of younger adults (Haier et al., 1988; also see Neabauer & Fink, 2009, for a review of the evidence) and that there is a strong correspondence between large-scale resting-state network connectivity based on fMRI and glucose network covariance in younger and older adults (Bernier et al., 2017; Di et al., 2012; Li et al., 2020; Tomasi et al., 2013, Tomasi et al., 2017), especially for hubs that have high metabolic rates (Liang et al., 2013; Dai et al., 2015).

Our second ALE analysis found a single area largely within the left thalamus with higher metabolic activity in older than younger adults, possibly reflecting a compensatory mechanism associated with ageing. Although this signal was the result of two studies only (n = 172 subjects; see tables 1 and 2), the increase metabolism in the thalamus in the second analysis (cluster 1) may reflect a compensatory mechanism for a decrease seen in the other regions in the first analysis (clusters 1 to 8), including other areas of the thalamus (cluster 9). The different pattern of metabolism in areas of the thalamus also highlights the potential for studies that only examine cerebral metabolism at the global or lobar level to miss potentially important signals that voxel-based analysis could identify. Additional research is needed to investigate the potential for metabolic compensatory affects both in the thalamus and other regions of the brain.

Although the results of the current meta-analyses do not directly test the compensation and efficiency hypotheses, when considered in conjunction with the wider cognitive ageing literature, they do not support the idea that older adults are more efficient in their use of energy to achieve the same performance as young individuals. Rather, they suggest that they have fewer absolute metabolic resources to perform the same tasks, which the wider literature indicates that they often perform at lower levels. Based on this finding, adults who maintain functioning into older adulthood are likely able to do so based on the capacity to recruit additional neural networks or draw on “reserve” rather than being able to rely on fewer and more efficient metabolic resources.

### 4.6 Limitations and Future Research

The studies reporting CRM_GLC_ effect sizes and peak activation differences in glucose metabolism by age did not generally include middle-aged individuals, which restricts the age differences that can be characterised across the adult lifespan. Specifically, cognitive and functional network changes often display a quadratic pattern, with an inflection point at approximately the third to fifth decade of life (e.g., Wei et al., 2018; Luo et al., 2020). There is some evidence to support a similar pattern for cerebral glucose metabolism. Pardo et al. (2007) reported glucose uptake values in the anterior cingulate, medial prefrontal and dorsomedial thalamus to display a negative quadratic relationship from the second to ninth decade of life, peaking around the age of 30 years. Further research is needed to establish whether or not the cerebral glucose changes found in the meta-analysis follow a similar trajectory from younger to older adult ages.

For the pooled effect size meta-analyses, a quantitative comparison of the age cohorts is not possible from the reported data (some reported mean age; others age ranges). However, of the 19 studies reporting age-group differences, 15 (79%) compared younger adults with an age range or mean age in the second to third decade of life compared to older adults with an age range or mean age in the sixth to ninth decade of life. Three studies (16%) split the samples into younger and older groups with the division at age 50 years (Blesa et al., 1997; Moeller et al., 1996; Yoshi et al., 1998). For the eight ALE studies, a combination of categorical (old versus young) and continuous adult age range samples were used. It is possible that cohort differences in age profiles are impacting the variability of the results found in our meta-analyses.

The samples on which the meta-analyses were based were from the U.S., Europe and Asia (see Table 1 and 2). Although some reported education level, the socioeconomic backgrounds of the participants is largely unknown. Most studies included a mix of male and female participants; however, separate data were not reported in sufficient quantity to allow for sex differences to be analysed. Additional research is needed to investigate the generalisability of the results to other geographical and cultural areas, as well as among diverse socioeconomic samples. Sex differences in age-related metabolic changes also require further research. Differential rates of changes are likely to be associated with sex differences in cognitive task performance and may contribute to sex differences in dementia (Li & Singh, 2014).

The current findings help to characterise cerebral hypometabolism in ageing. The question remains whether metabolic differences observed in FDG-PET studies are reflecting isolated neuronal alterations in metabolism in ageing, or whether other factors are driving or moderating the age-related differences. For example, there is an increased risk of cognitive decline and dementia related to hypoglycaemia, glycaemic control, metabolic syndrome and insulin resistance (Akintola & van Heemst, 2015; Arvanitakis et al., 2016; Bello-Chavolla et al., 2019; Ekblad et al., 2017). It is also believed that these metabolic states act as mediators, moderators or even accelerators of age-related cognitive decline (Raz & Rodrigue, 2006). Given the rising rates of chronic diseases that impact metabolism in the periphery and brain, such as obesity (Afshin et al., 2017) and diabetes (Tao et al., 2015), it is likely that future research in cognitive ageing may find an important role of peripheral factors, particularly as age itself is a key risk factor for these diseases. Better understanding the influence of metabolic states on cognitive ageing may also allow them to be used as early markers for age-related cerebral metabolic and associate cognitive changes.

The pooled effect size meta-analyses drew on studies that used several different approaches to adjust for brain volume changes with age and control for partial volume effects. The approaches included co-registration to CT scans (e.g., de Leon 1983, 1984) and MRI (e.g., Blesa et al., 1987), controlling for ventricle size (Kushner et al., 1987) and cerebrospinal fluid (Chawluk et al., 1990), and partial volume effect correction with MRI coregistration (Nugent et al., 2014a, b). The heterogeneity was higher in the corrected compared to the non-corrected samples (except for the frontal lobe), suggesting that the different approaches either have variable effects in adjusting for volume changes with age or are too heterogeneous to show age differences with confidence among the relatively small number of samples available. Only a subset of the corrected samples used partial volume effect correction, also raising the possibility that measurement error accounts for at least some of the age group differences found in the overall and uncorrected analyses. For the ALE analyses, we had insufficient samples to undertake contrast analysis between PVE-corrected and uncorrected samples. Hence, we were limited to an exploratory, non-quantitative comparison. Further research is needed to assess whether age-related changes in metabolism are sensitive to the different measures of brain volume changes. In line with previous authors (e.g., Metzer et al., 1999; Greve et al., 2016), we recommend that partial volume effect correction be a standard in future quantitative PET research in metabolic ageing.

It is well established from fMRI research that brain activity at rest and during task performance is organised into functional networks, defined by their spatiotemporal configuration and functional roles (Beckmann et al., 2005; Calhoun et al., 2008). The use of FDG-PET to study brain network has historically been limited by poor temporal resolution, with studies acquiring single images that index the rate of glucose uptake across the entire scan. To be comparable to fMRI functional network connectivity measures, it is necessary to measure the time-course of FDG-PET activity across the scan for each individual subject, and then correlate those time-courses across regions to form a connectome. Recent advances in FDG-PET data acquisition have made it possible to measure the time course of FDG uptake over the course of a scan (Villien et al., 2014; Hahn et al., 2016; Jamadar et al., 2019). This ‘fPET’ method has been applied to characterise metabolic connectivity in younger adults (Li et al., 2020; Jamadar et al., 2020; Jamadar et al., 2021) and offers promise for expanding metabolic network analysis in ageing.

## 5. Conclusion

The results of the systematic review and meta-analyses suggest that there is both a global and lobular reduction in the absolute rate of cerebral glucose metabolism between the third and sixth decades of life. The largest and most robust contribution comes from the frontal lobe, with changes also seen in the temporal lobe and accompanied by smaller regions of hypometabolism. The results also suggest that adjusting for brain volume changes and partial volume effects should be standard practice in studies of brain metabolic differences in ageing. Ageing related metabolic changes in frontal and temporal regions and in selected sub-cortical structures are important for interpreting theories of cognitive ageing and in distinguishing “normal” ageing from neurodegenerative diseases. Further elucidating the mechanisms driving reduced glucose metabolism and their impact on cognitive ageing is an important empirical and clinical matter, as they may serve as early markers or targets for interventions to slow or prevent age-related related decline.

## Data Statement

The data used in this paper are publicly available from published journals articles.

## Author contributions

HD and SJ designed the review. HD and RD undertook the review of database searches, reconciliation of included papers, independent entry and counts of the data for the meta-analyses. HD wrote the first draft of the review with SJ edits. All authors contributed to manuscript preparation or review.

## Supplementary Material

### No Significant Effect of ROI Averaging

We undertook additional analyses on the CRM_GLC_ effect sizes to investigate the potential impact of averaging regions that were not always of the same size. Two of the five studies that did not report CRM_GLC_ values for a lobe(s) reported grey matter ROI volumes or referenced templates for which volumes could be obtained (Nugent et al., 2014a,b). Volumes were not available for the remaining three studies (Azari et al., 1992; Eberling et al., 1995; Moeller et al., 1996). We first excluded all five studies and repeated the pooled effect size analyses. We also repeated the analyses including two studies and weighting the ROIs by their relative volume within the lobe (Nugent et al., 2014a.b), and excluding the remaining three studies for which volume data were not available.

The analyses excluding the five studies reporting ROI CRM_GLC_ within lobes, as well as the analyses including the two studies for which the ROIs were weighted by their volume, are available in the Supplementary Materials (Tables S1 and S2). Both sets of analyses yield results consistent with those reported in section 3.1 and 3.2. Specifically, the strongest age-related metabolic reduction was found in the frontal lobe and it remained significant after volume correction (excluding all five studies, confidence interval -1.06, to -0.15; retaining and weighting two studies, confidence interval -0.90 to -0.33). Moreover, the corrected vs uncorrected results across the other lobes are consistent with the results reported for all samples.

### Contrast Analysis of PVE-corrected and Uncorrected Samples

We undertook contrast analysis in GingerALE to test the PVE-corrected and uncorrected sets of foci for statistically significant differences in spatial convergence. This allowed us to localise foci commonly or differentially activated in the PVE-corrected and uncorrected samples. Contrast analysis compares two ALE datasets using the voxel-wise minimum value of the input ALE images (Eickhoff et al., 2012; Laird et al., 2005). An image for each dataset is created, then one is subtracted from the other and compared to the combined data. GingerALE generates a conjunction image that shows the similarity between the datasets. Hence, we undertook separate ALE analyses on the PVE-corrected and uncorrected samples (n = 4 and 7, respectively; see Table 2) as input into the contrast analysis in GingerALE. We also overlaid each map onto a composite image to visualise the locations of conjunction and contrast in the ALE clusters

Results for the contrast analyses comparing the PVE-corrected and uncorrected samples showed three areas of conjunction in the inferior and medial frontal gyri, overlapping clusters 1, 2 and 4 from the analysis of the combined samples (see Table S3 and S4 and Figure S1A-C). Separate locations were found in the anterior cingulate in the PVE-corrected and uncorrected samples (Figure S1D). The locations were spatially close although not statistically significant in the contrast analysis. A cluster from PVE-corrected samples only was identified in the superior temporal lobe, equating to cluster 7 in combined analysis (Figure S1E). Clusters from the uncorrected samples only were found in the cingulate, equating to cluster 8 in the combined analysis (Figure S1E); and a second cluster in the caudate, corresponding to cluster 5 in the combined samples (Figure S1F).

**Figure S1.**
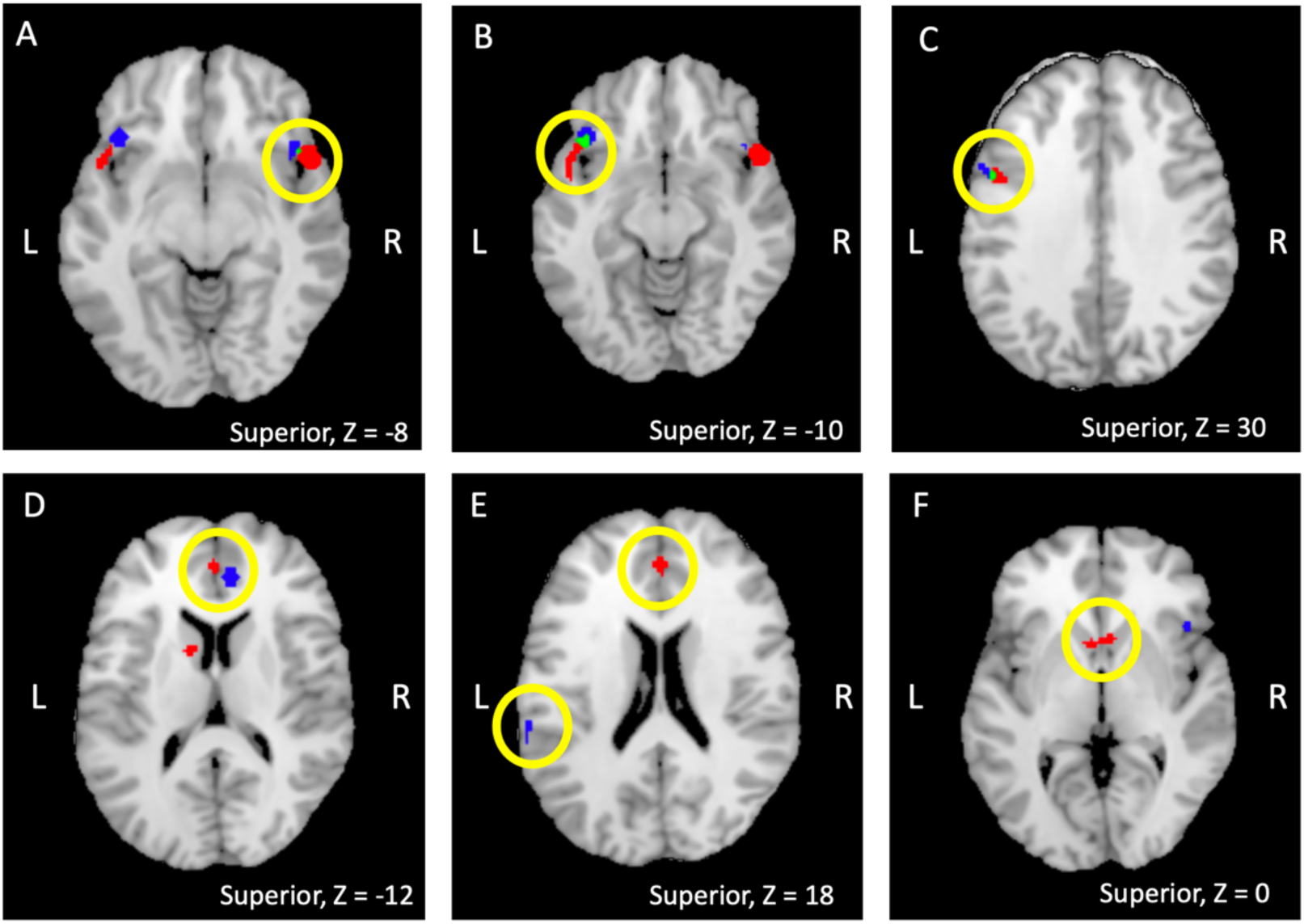
Statistical comparisons of ALE maps. ALE meta-analyses were carried out separately for samples with PVE correction (blue) and without PVE correction (red) and are presented with the overlays from the separate meta-analyses. Green indicates overlap between the individual meta-analyses from the conjunction analysis. Panels A, B and C show areas of conjunction between the two sets of samples and approximate clusters 1, 2 and 4 from the analysis of all combined samples. Panel D shows separate location in the anterior cingulate from the corrected and non-corrected samples, approximating cluster 3 in the combined analysis but not identified in the conjunction analysis. Panel E showing a cluster from PVE corrected sample only in the superior temporal lobe (blue), equating to cluster 7 in combined analysis; and a cluster from non-corrected samples only in the cingulate (red), equating to cluster 8 in the combined analysis. Panel F showing cluster from non-corrected samples only in the caudate, corresponding to cluster 5 in the analysis of the combined samples.

**Table S1.** Summary of meta-analyses findings. For Hedges’ g, numbers in parentheses are 95% confidence intervals.

|  | Samples | N-Size | Mean % Reduction<br>Younger vs older | Hedge's G<br>Effect Size | Z-Value | Heterogeneity Q | Heterogeneity (I<sup>2</sup>) |
| --- | --- | --- | --- | --- | --- | --- | --- |
| **Whole brain: Correlation with Age** | 13 | 567 |  | -0.24 (-0.33, -0.14) | -5.22, p = .001 | 13.9, p = .305 | 13.8% |
| **Whole brain: Age Group differences** |  |  |  |  |  |  |  |
| All samples (inc averaged ROIs) | 15 | 620 | 7% | -0.47 (-0.74, -0.20) | -3.73, p = .000 | 25.9, p = .027 | 45.9% |
| Excluding samples reporting ROIs only | 13 | 566 | 7% | -0.49 (-0.77, -0.22) | -3.88, p = .000 | 20.58, p = .013 | 41.7% |
| **Frontal** |  |  |  |  |  |  |  |
| All samples (inc averaged ROIs) | 13 | 561 | 12% | -0.66 (-0.89, -0.43) | -6.29, p = .000 | 15.1, p = .238 | 20.3% |
| Excluding samples reporting ROIs only | 8 | 275 | 15% | -0.73 (-1.15, -0.32) | -4.17, p = .000 | 11.9, p = .104 | 41.2% |
| Retained and weighted ROI sampis by volume where possible | 10 | 379 | 13% | -0.70 (-1.01, -0.42) | -5.46, p < .000 | 12.00, p = .215 | 24.9% |
| **Temporal** |  |  |  |  |  |  |  |
| All samples (inc averaged ROIs) | 12 | 546 | 11% | -0.56 (-0.90, -0.22) | -3.67, p = .000 | 24.5, p = .011 | 55.0% |
| Excluding samples reporting ROIs only | 6 | 243 | 11% | -0.51 (-1.18, 0.17) | -1.93, p = .027 | 15.43, p = .009 | 67.6% |
| Retained and weighted ROI sampis by volume where possible | 8 | 347 | 10% | -0.48 (-0.92, -0.04) | -2.56, p = .005 | 16.28, p = .023 | 57.0% |
| **Parietal** |  |  |  |  |  |  |  |
| All samples (inc averaged ROIs) | 13 | 559 | 9% | -0.37 (-0.72, -0.02) | -2.30, p = .011 | 29.9, p = .003 | 59.8% |
| Excluding samples reporting ROIs only | 8 | 273 | 11% | -0.34 (-0.98, 0.30) | -1.24, p = .107 | 26.08, p = .000 | 73.2% |
| Retained and weighted ROI sampis by volume where possible | 10 | 377 | 9% | -0.30 (-0.77, -0.17) | -1.43, p = .076 | 26.25, p = .002 | 65.7% |
| **Occipital** |  |  |  |  |  |  |  |
| All samples (inc averaged ROIs) | 8 | 414 | 8% | -0.39 (-0.64, -0.14) | -3.63, p = .000 | 7.6, p = .370 | 7.8% |
| Excluding samples reporting only ROIs | 3 | 128 | 15% | -0.71 (-1.92, 0.49) | -2.55, p = .005 | 3.74, p = .154 | 46.6% |
| Retained and weighted ROI sampis by volume where possible | 7 | 382 | 9% | -0.40 (-0.70, -0.11) | -3.40, p < .000 | 6.97, p = .324 | 13.9% |
| **White Matter: Age Group differences** | 6 | 265 | 3% | -0.45 (-1.90, 1.00) | -0.80, p = .213 | 50.2, p < .001 | 90.4% |
| **Subcortical: Age Group differences** |  |  |  |  |  |  |  |
| Caudate | 9 | 400 | 9% | -0.50 (-0.89, -0.10) | -2.89, p = .002 | 20.3, p = .009 | 60.7% |
| Thalamus | 9 | 400 | 8% | -0.40 (-0.85, 0.05) | -2.05, p = .020 | 23.0, p = .003 | 65.2% |
| Cerebellum | 5 | 243 | 3% | -0.20 (-1.36, 0.96) | -0.48, p = .317 | 42.4, p < .001 | 90.6% |

**Table S2.**
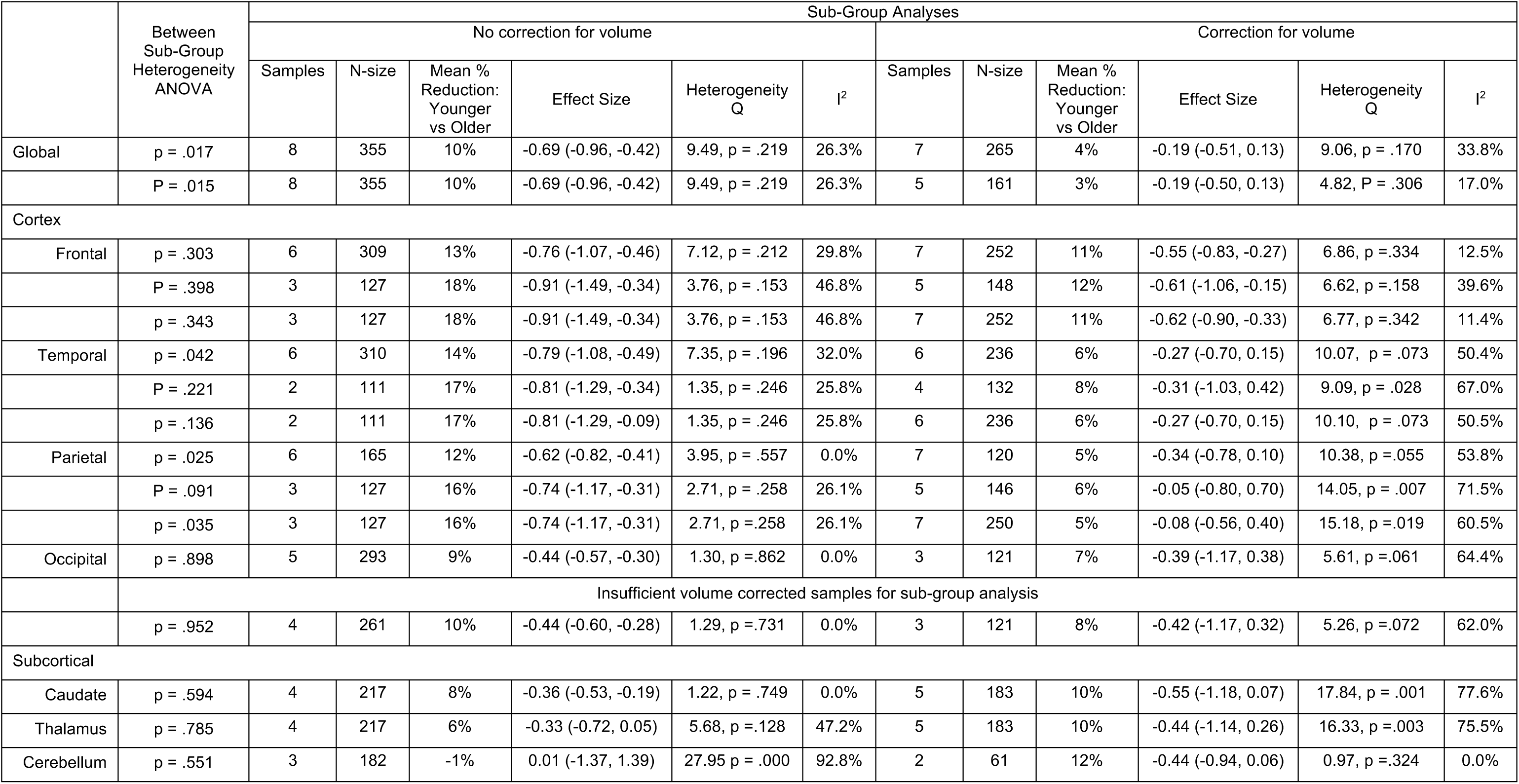
Sub-group analyses for studies with and without brain volume correction in ageing. Numbers in parentheses are 95% confidence intervals. For each lobe, top row is all samples (including averaged ROIs) from Table 3 in main paper; middle excludes all samples reporting only ROIs that were averaged in the top row; and bottom row excludes all samples reporting only ROIs that were averaged where volume adjustments could not be made but includes samples were volume adjustments were possible.

**Table S3.** Conjunction analysis of locations (Talairach) of overlap in ALE clusters of PVE-corrected and non-corrected samples.

| Cluster Number | Size (mm<sup>3</sup>) | Hemisphere | Lobe | Gyrus | Brodmann's Area(s) | Peak Coordinate X, Y, Z | ALE Value |
| --- | --- | --- | --- | --- | --- | --- | --- |
| 1 | 120 | Left, 100% | Frontal, 100% | Inferior frontal, 100% | 47 (100%) | -40, 20, -12 | 0.0107 |
| 2 | 88 | Left, 100% | Frontal, 100%<br>Frontal, 48.4% | Inferior frontal , 71%<br>Middle frontal, 29% | 9 (100%) | -48, 6, 32 | 0.0107 |
| 3 | 16 | Right, 100% | Frontal, 100% | Inferior frontal, 100% | 47 (100%) | 44, 16, -7 | 0.0088 |

**Table S4.** Locations (Talairach) of significant clusters from the ALE meta-analysis of glucose metabolism differences for older vs younger adults for PVE-corrected and uncorrected samples.

| Cluster Number | Size (mm<sup>3</sup>) | Hemisphere | Lobe | Gyrus | Brodmann's Area(s) | No of Samples (sample reference No., Table 2) | Peak Coordinate X, Y, Z | ALE Z | ALE Value |
| --- | --- | --- | --- | --- | --- | --- | --- | --- | --- |
| Clusters of **lower** glucose metabolism among older versus younger adults: PVE corrected samples |  |  |  |  |  |  |  |  |  |
| 1 | 568 | Left, 100% | Frontal, 100% | Inferior frontal, 100% | 47 (100%) | 2 (9, 10) | -38, 22, -10 | 5.13 | 0.0176 |
| 2 | 496 | Right, 100% | Sub-lobar, 52%<br>Frontal, 48.4% | Inferior frontal, 48%<br>Insula, 48%<br>Extra-nuclear, 3% | 47 (55%)<br>13 (36%) | 2 (2, 10) | 42, 18, -6 | 4.13 | 0.0114 |
| 3 | 440 | Right, 100% | Limbic, 100% | Anterior cingulate, 100% | 32 (94%) | 2 (9, 10) | 8, 38, 12 | 4.64 | 0.0151 |
| 4 | 200 | Left, 100% | Temporal, 100% | Superior temporal, 100% | 22 (60%)<br>42 (40%) | 2 (1, 2) | -60, -32, 20 | 3.74 | 0.0098 |
| 5 | 200 | Left, 100% | Frontal, 100% | Inferior frontal, 81%<br>Middle frontal, 19% | 9 (100%) | 2 (1,10) | -46, 6, 32 | 4.05 | 0.0107 |
| Clusters of **lower** glucose metabolism among older versus younger adults: Uncorrected samples |  |  |  |  |  |  |  |  |  |
| 1 | 1,128 | Right, 100% | Temporal, 69%<br>Frontal, 31% | Superior temporal, 69%<br>Inferior frontal, 31% | 38 (51%)<br>47 (31%)<br>22 (19%) | 4 (3, 4, 8, 11) | 50, 12, -14 | 6.29 | 0.0276 |
| 2 | 864 | Left, 82%<br>Right, 18% | Sub-lobar, 99%<br>Limbic, 1% | Caudate, 99%<br>Anterior cingulate, 1% |  | 4 (3, 4, 6, 8) | -8, 8, 4 | 3.85 | 0.0122 |
| 3 | 824 | Left, 100% | Temporal, 57%<br>Frontal, 43% | Superior temporal, 49%<br>Inferior frontal, 43%<br>Middle temporal, 8% | 38 (43%)<br>47 (43%)<br>21 (8%)<br>22 (5%) | 4 (4, 6, 7, 8) | -48, 4, -14 | 3.91 | 0.0126 |
| 4 | 424 | Right, 50%<br>Left, 50% | Limbic, 50%<br>Frontal, 50% | Anterior cingulate, 50%<br>Medial frontal, 50% | 32 (50%)<br>9 (50%) | 2 (4, 8) | 0, 42, 16 | 4.31 | 0.0152 |
| 5 | 312 | Left, 100% | Frontal, 100% | Inferior frontal, 62%<br>Middle frontal, 24%<br>Precentral, 14% | 9 (91%)<br>6 (9%) | 2 (5, 6) | -46, 6, 32 | 4.19 | 0.0146 |
| 6 | 288 | Right, 100% | Frontal, 61% | Cingulate, 100% | 32 (100%) | 2 (4, 8) | 4, 20, 36 | 3.93 | 0.0128 |

